# SPRTN protease and SUMOylation coordinate DNA-protein crosslink repair to prevent genome instability

**DOI:** 10.1101/2020.02.14.949289

**Authors:** Bruno Vaz, Annamaria Ruggiano, Marta Popovic, Gonzalo Rodriguez-Berriguete, Susan Kilgas, Abhay N. Singh, Geoffrey S. Higgins, Anne E. Kiltie, Kristijan Ramadan

**Author notes:** these authors contributed equally (Bruno Vaz, Annamaria Ruggiano). leading contact (Kristijan Ramadan). Ruđer Bošković Institute, Bijenička cesta 54, 10000 Zagreb, Croatia (Marta Popovic).

## Abstract

DNA-protein crosslinks (DPCs) are a specific type of DNA lesions where proteins are covalently attached to DNA. Unrepaired DPCs lead to genomic instability, cancer, neurodegeneration and accelerated ageing. DPC proteolysis was recently discovered as a specialised pathway for DPC repair. The DNA-dependent SPRTN protease and 26S proteasome emerged as as two independent proteolytic systems for DPC repair. DPCs are also repaired by homologous recombination (HR), a canonical DNA repair pathway. While studying the role of ubiquitin and SUMO in DPC repair, we identified mutually exclusive signalling mechanisms associated with DPC repair pathway choice. DPC modification by SUMO-1 favours SPRTN proteolysis as the preferred pathway for DPC repair. DPC SUMOylation counteracts DPC ubiquitination, which promotes DNA breaks and the switch to HR. We propose that modification of DPCs by SUMO-1 promotes SPRTN proteolysis, which is essential for DPC removal to prevent DNA replication defects, chromosomal recombination and genomic instability.

## INTRODUCTION

DNA-protein crosslinks (DPCs) are ubiquitous and heterogeneous DNA lesions that arise from irreversible, covalent binding of a protein to DNA following exposure to a chemical or physical crosslinking agent, *e.g.* formaldehyde or UV light (Fielden et al., 2018; Ide et al., 2018; Vaz et al., 2017). Formaldehyde is a cellular by-product of methanol metabolism, histone demethylation and lipid peroxidation as well as an environmental pollutant. It is estimated that intracellular formaldehyde concentrations can reach 400 µM (Andersen et al., 2010), implying that threats posed by DPC are ubiquitous. Furthermore, some of the most commonly used chemotherapeutics, namely the topoisomerases (Topo) 1 and 2 poisons camptothecin (CPT) and etoposide cause abortive topoisomerase activity on DNA; this then causes a specific class of DPCs known as Topo-1 or -2 cleavage complexes (Topo-1/2-ccs) (Ashour et al., 2015; Pommier and Marchand, 2012). Due to the stability of the crosslink and their bulkiness, DPCs constitute a barrier to all DNA transactions. If left unrepaired, DPCs lead to genomic instability and/or cell death as well as disease conditions including neurodegeneration, cancer and premature ageing in humans and mice (Gómez-Herreros et al., 2014; Lessel et al., 2014; Maskey et al., 2014, 2017). To cope with DPC-induced toxicity, cells employ two major DPC repair mechanisms: (i) a proteolytic-dependent mechanism, where the proteinaceous part of the DPC is cleaved by specific proteases during replication, and (ii) a nucleolytic-dependent mechanism, where the nucleases involved in homologous recombination (HR) or nucleotide excision repair cleave off DNA bearing a crosslinked protein (Aparicio et al., 2016; Hoa et al., 2016; Nakano et al., 2007, 2009). The former mechanism involves DNA-dependent metalloproteases, SPRTN in metazoans and Wss1 in yeast (Lopez-Mosqueda et al., 2016; Maskey et al., 2017; Mórocz et al., 2017; Stingele et al., 2014, 2016; Vaz et al., 2016), and the proteasome (Larsen et al., 2019; Sparks et al., 2019). A putative protease, acid repeat containing protein (ACRC), also known as germ cell nuclear antigen (GCNA), has recently been discovered and linked to DPC repair (Borgermann et al., 2019; Fielden et al., 2018). While both proteolytic and nucleolytic pathways protect cells from DPC-induced toxicity, they come with downsides. HR can lead to aberrant genomic rearrangements and loss of heterozygosity (Liu et al., 2012), while proteolytic pathways can increase mutagenesis (Mórocz et al., 2017; Stingele et al., 2014). However, it is not known how DPC pathway choice between proteolysis and HR is coordinated.

Recently, post-translational modifications on DPCs by ubiquitin (Ub) or small Ub-like modifier (SUMO) molecules have emerged as two signals that govern proteolysis-dependent DPC repair: ubiquitination of DPCs promotes the 26S proteasome proteolysis repair during DNA replication in *Xenopus* egg extract, while SUMOylation of DPCs promotes ACRC recruitment to DPC lesions and their repair outside DNA replication (Borgermann et al., 2019; Larsen et al., 2019; Sparks et al., 2019). Considering that ACRC is expressed predominantly in germ and stem cells and not in human primary or cancer cell lines, it remains unclear why SUMOylation is important for DPC repair in proliferative somatic human cells.

Here, we report that both ubiquitin and SUMO are required for SPRTN-dependent DPC repair during replication. We show that SPRTN clears SUMOylated DPCs in an ubiquitin-dependent manner. In parallel, we show that SUMOylation suppresses HR-mediated recombinogenic events. Simultaneous inactivation of SPRTN-dependent proteolysis and HR leads to synthetic lethality after formaldehyde exposure, suggesting that SPRTN and HR act in parallel and not within the same DPC repair pathway to prevent DPC-induced toxicity. We propose that SUMOylation channels DPC pathway choice towards SPRTN-dependent proteolysis to prevent recombinogenic events that could lead to genomic instability.

## RESULTS

### DPC-inducing agents trigger SUMOylation and ubiquitination waves

To gain insights into the ubiquitin and SUMO signals associated with DPC repair in proliferative mammalian cells, we analysed the dynamics of both posttranslational modifications (PTMs) on DPCs or soluble protein fraction following exposure to the general DPC-inducing agent formaldehyde (FA) (Figure 1A, 1B and S1A). DPCs were rapidly formed upon 10 minutes’ pulse with FA in a dose-dependent manner. DPCs underwent extensive modification by SUMO-1, SUMO-2/3 and ubiquitin (Figure 1B, right panels). Similar ubiquitination and SUMOylation patterns were observed in the soluble protein fraction (Figure S1A). Longer incubation times with FA did not increase DPCs but rather decrease their amount, suggesting the activation of fast DPC repair mechanisms (Figure S1B). FA can also form interstrand crosslinks (ICLs), which are resolved by the Fanconi anemia pathway (Ceccaldi et al., 2016). To rule out its involvement in our experimental setup, we monitored the ubiquitination status of FANCD2, a recognised marker for activation of the Fanconi anemia pathway. In contrast to mitomycin C (MMC) and cisplatin (cis) treatment, known ICL-inducing agents, FA treatment did not induce mono-ubiquitination of FAND2 (Figure S1C, upper panel). Moreover, MMC or cisplatin neither increased SUMOylation nor induced DPCs (Figure S1C lower panel, and S1D), further indicating that our experimental setup with FA highlighted ubiquitination and SUMOylation signals specifically associated with DPC and not ICL induction. To confirm the observed PTM signature on DPCs, we analysed the PTM pattern following treatment with the Topo-1 inhibitor camptothecin (CPT). CPT treatment specifically trapped Topo-1 in a dose-dependent manner (Figure S1E), and, similarly to FA, induced ubiquitination and SUMOylation in DPCs and soluble fractions (Figure S1F). The specificity of CPT for Topo-1ccs formation accounted for the fainter PTM signal observed in the DPC fraction (Figure S1F, right panel) than observed after FA treatment (Fig. 1B).

**Figure 1.**
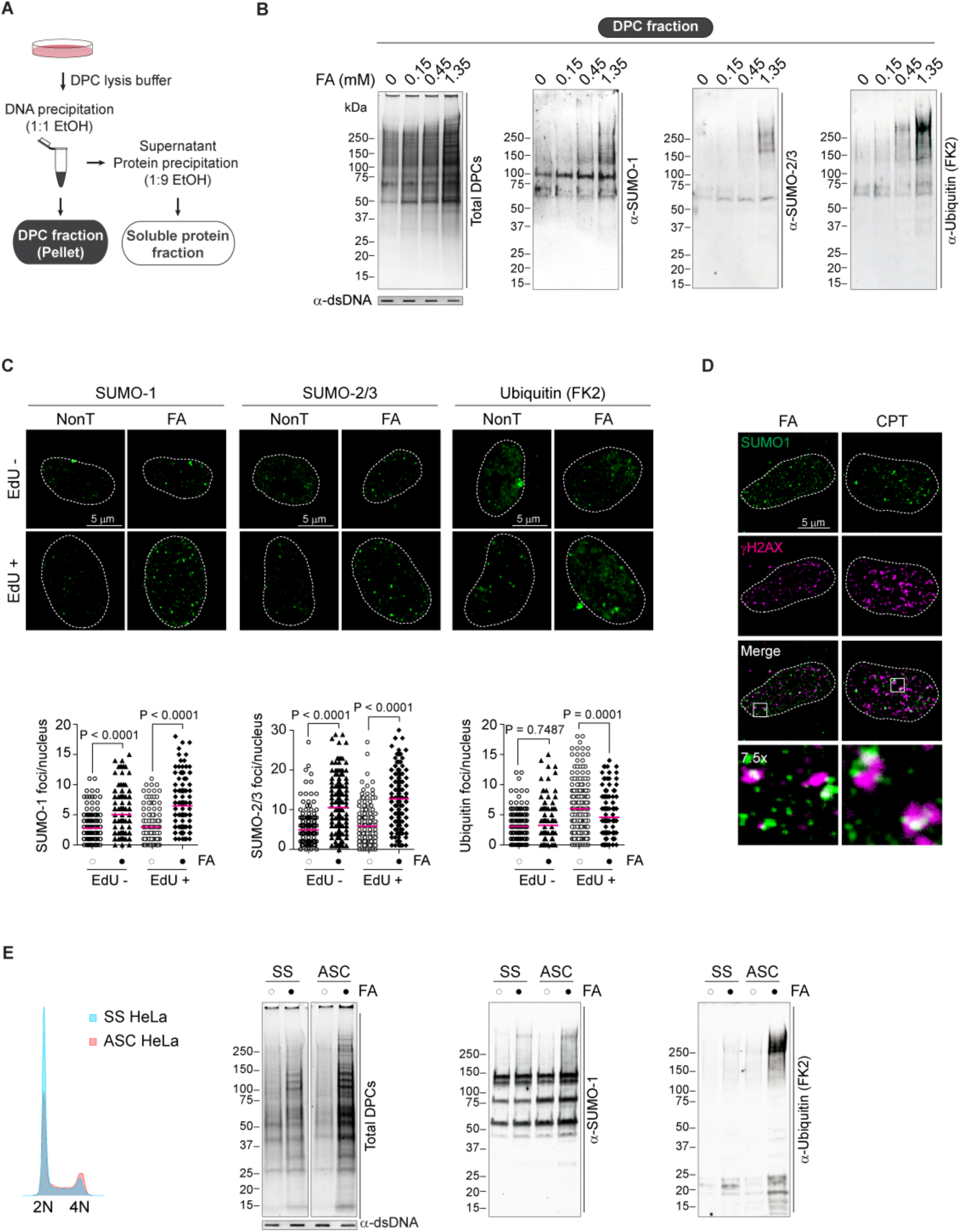
**DPC-inducing agents trigger SUMOylation and ubiquitination waves** (A) Schematic of the DPC and soluble fraction isolation protocol. (B) FA treatment promotes ubiquitination and SUMOylation on DPCs. HeLa cells were treated with increasing concentrations of formaldehyde (FA) for 10 minutes at 37°C and fractions were isolated as in A. DPCs were analysed by western blot for the indicated post-translational modifications (PTMs). Double stranded (ds) DNA was used as a loading control that DPCs were isolated from the same amount of genomic DNA. (C) FA treatment causes SUMO foci formation. RPE-1 cells were treated with 1 mM FA for 10 minutes at 37°C. EdU was added 20 minutes before FA treatment in order to label dividing cells. After treatment, cells were pre-extracted, fixed and immunostained with the indicated antibodies. Bottom panel indicates quantifications of the number of foci per nucleus. Foci were counted with ImageJ (200 nuclei) and statistical significance calculated using unpaired t-test. (D) DPC-induced SUMO-1 foci partially co-localise with γH2AX. RPE-1 cells were treated with 1 mM FA for 10 minutes at 37°C. After treatment, cells were pre-extracted, fixed and immunostained with the indicated antibodies. (E) DPCs form mostly in cycling cells. Asynchronous (ASC) and serum-starved (SS) HeLa cells were treated with 1.35 mM FA for 10 minutes at 37°C. Total DPCs were isolated by RADAR and detected with Flamingo protein gel staining (left panel) or analysed by western blot for the indicated PTMs. FACS (far left panel) analysis of the DNA content (propidium iodide) confirms a partial cell cycle arrest of SS cells.

We proceeded to test whether FA induced accumulation of these PTMs in nuclear foci. Short FA pulses lead to a 2-fold increase in the average number of SUMO-1 and SUMO-2/3 foci both in S-phase (EdU-positive) and non-S-phase (EdU-negative) RPE-1 cells (Figure 1C). Unlike SUMO, ubiquitin did not accumulate at specific foci but rather in a pan-nuclear pattern (Figure 1C and S1G). Confocal microscopy showed a partial co-localisation of the SUMO-1 foci with the general DNA damage marker γH2AX, suggesting that, at least in part, SUMO-1 accumulates at FA- and CPT-induced DNA damage sites (Figure 1D).

To address the association between cell proliferation and FA-induced DPCs, we analysed DPC formation in replicative (asynchronous) versus G0-arrested cells using two models: (1) HeLa cells in which we induced G1/G0-like state by serum starvation, and (2) RPE-1 cells, which arrest at G0 when confluent. Analysis of DNA content by flow cytometry confirmed a partial and total arrest in G1/G0, in HeLa and RPE-1 cells, respectively (Figure 1E and S1H). FA mostly triggered DPC formation in asynchronous proliferative cells and these DPCs were SUMOylated (SUMO-1) and ubiquitinated (Figure 1E and S1H). When we blocked DNA replication with hydroxyurea (HU) and treated the cells with FA, DPC SUMOylation and ubiquitination increased, suggesting that DNA replication is involved in removal of modified DPCs (Figure S1I). We conclude that DPC-inducing agents trigger rapid ubiquitination and SUMOylation waves, which are removed in a DNA replication-dependent manner.

### SPRTN prevents accumulation of SUMOylated DPCs

We and others identified SPRTN as a DNA-dependent metalloprotease required for DPC proteolysis during DNA replication (Lopez-Mosqueda et al., 2016; Maskey et al., 2017; Mórocz et al., 2017; Stingele et al., 2016; Vaz et al., 2016). To investigate SPRTN’s role in the proteolysis of modified DPCs, we analysed the ubiquitin and SUMO signals following recovery from FA in parental and SPRTN haplo-insufficient (ΔSPRTN) HeLa cells. As previously reported, DPC removal was significantly affected in SPRTN-deficient cells (Figure S2A) (Vaz et al., 2016). In parental cells modified DPCs were removed within 20 minutes of recovery from FA treatment. In contrast, SPRTN-deficiency caused delayed repair of SUMOylated DPCs (SUMO-1), while ubiquitinated DPCs were removed within 60 min (Figure 2A and S2B). DPC removal defects in ΔSPRTN cells were not caused by cell cycle defects or replication arrest (Figure S2C and S2D). In contrast to the DPC fraction, SPRTN depletion had lesser or no effect on removal of the SUMO and ubiquitin signals in the soluble protein fraction (Figure S2E). To further test the involvement of SPRTN in the proteolysis of modified DPCs, we monitored DPC modifications following synchronous progression through S-phase (Figure S3A and S3B). SPRTN-depleted cells, unlike parental cells, failed to process DPCs during S-phase progression (Figure S3A) (Vaz et al., 2016). While the ubiquitin signal was partially cleared from the DPCs, SUMO-1 persisted (Figure S3B). Together, these results indicate that SPRTN prevents accumulation of SUMO-1-DPCs.

**Figure 2.**
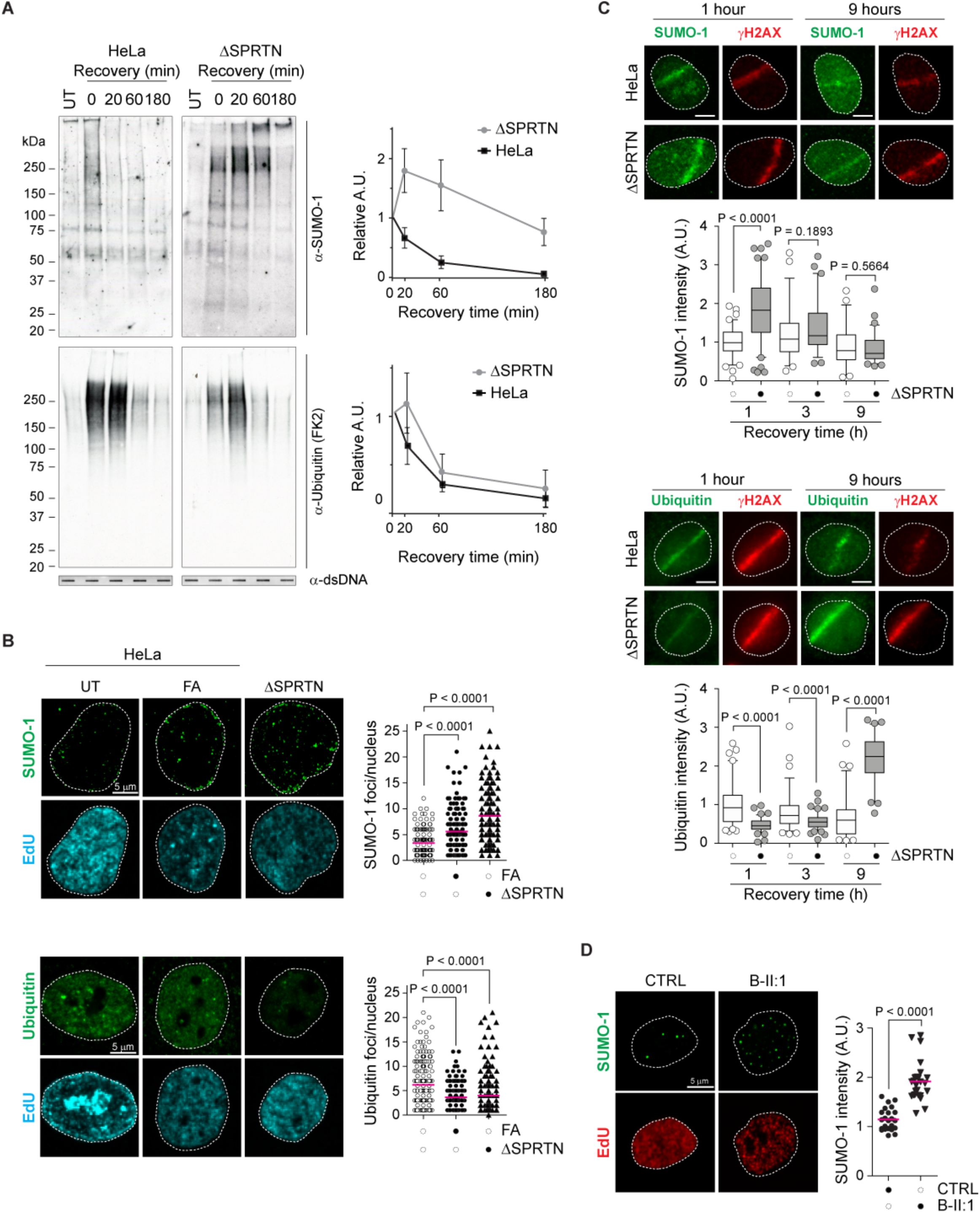
**SPRTN prevents accumulation of SUMOylated DPCs** (A) SPRTN-deficient cells are unable to process SUMO-DPCs. Parental and ΔSPRTN HeLa cells were treated with 1.35 mM FA for 10 minutes at 37°C and allowed to recover for the indicated times. Total DPCs were isolated by RADAR and analysed by western blot for the indicated PTMs. Graphs show the mean ± SEM of relative signal from 3 independent experiments. (B) SPRTN prevents accumulation of SUMO-1 foci. Wild-type Hela cells were treated with 1.35 mM FA for 10 minutes at 37°C. No FA was added to ΔSPRTN HeLa cells. Cells were pre-extracted, fixed and immunostained with the indicated antibodies. Foci were counted from EdU positive cells (200 nuclei) with ImageJ and statistical significance calculated using unpaired t-test. (C) SPRTN-deficient cells accumulate SUMO-1 at DNA damage sites induced by UV laser microirradiation. Wild-type and ΔSPRTN HeLa cells were sensitized with Hoechst 33258 for 30 minutes prior to laser microirradiation. EdU was added during the sensitization time. Cells were allowed to recover from damage for the indicated times. Cells were then pre-extracted, fixed and immunostained with the indicated antibodies. Signal at DNA damage sites was quantified in EdU positive cells (100 stripes) using ImageJ and statistical significance calculated using unpaired t-test. (D) RJALS patient cells accumulate SUMO-1 foci. Normal MRC5 and patient B-II:1 primary fibroblasts were pre-extracted, fixed and immunostained with the indicated antibodies. SUMO-1 signal was quantified in EdU positive cells (25 nuclei) using ImageJ and statistical significance calculated using unpaired t-test.

In addition, SPRTN-deficient cells (ΔSPRTN or siRNA) accumulated SUMO-1 foci (Figure 2B). Foci partially co-localised with γH2AX, suggesting their association with damage sites (Figure S3C). In contrast, no ubiquitin foci were detected in SPRTN-deficient cells (Figure 2B). In fact, nuclear ubiquitin staining was weaker compared to control cells (Figure 2B, right bottom panel), thus hinting a possible competition between SUMO and ubiquitin signals. To explore this, we analysed these PTMs at DNA damage sites induced by UV laser micro-irradiation (Figure 2C) (Davis et al., 2012; Stingele et al., 2016). ΔSPRTN cells showed a 2-fold increase in the SUMO-1 signal at damage sites following 1-hour recovery. On the contrary, ubiquitin signal decreased by 2-fold. A similar albeit less pronounced effect was observed 3 hours after UV laser micro-irradiation (Figure 2C). Interestingly, at 9 hours post-irradiation, a drop in the SUMO-1 signal and a robust increase in the ubiquitin signal were detected in ΔSPRTN cells. Altogether, these results confirm the role of SPRTN in preventing hyper-accumulation of SUMO-1 conjugates at DNA damage sites. Moreover, they suggest that SUMO-1 counteracts ubiquitin accumulation in SPRTN-deficient cells in the first several hours after DNA damage, an observation that will be elaborated upon later in this work.

Lastly, to address the relevance of the SPRTN-SUMO axis in human pathogenesis, we monitored SUMOylation in SPRTN-deficient cells from a Ruijs-Aalfs syndrome (RJALS) patient (Lessel et al., 2014). RJALS primary fibroblasts showed higher SUMO-1 intensity and foci number compared to control fibroblasts (Figure 2D). Similarly, we observed an accumulation of SUMO-1 conjugates in the chromatin of a patient’s lymphoblastoid cell line following a short pulse with FA (Figure S3D).

Altogether, these results indicate that SPRTN prevents accumulation of SUMOylated DPCs.

### SPRTN clears SUMOylated proteins

To investigate whether SPRTN processes SUMO and Ub modified substrates we took advantage of the “trapping” effect of inactive protease mutants (Flynn et al., 2003; Westphal et al., 2012). We immunopurified Flag-tagged SPRTN-WT or -E112A from HEK293 cells under physiological conditions (150 mM NaCl) (Figure 3A and S4A). SUMO-1, SUMO2/3 and ubiquitin were detected in SPRTN immunoprecipitates. However, only SUMO-1 and ubiquitin, but not SUMO-2/3, -modified substrates increased in SPRTN^E112A^ pull-down compared to SPRTN^WT^ (Figure 3A and S4A). To rule out the possibility that direct modifications on SPRTN could account for the PTM signal, we incubated the immunopurified SPRTN in high stringency buffer (1 M NaCl) (Figure S4B). Under these conditions, we observed that most of SPRTN’s interactions with SUMO-1 and ubiquitin, as well as with a known SPRTN interacting partner, the ATPase p97, were lost. These results indicate that SPRTN interacts with SUMO-1 and ubiquitin substrates in cells. To complement these results we analysed co-localization of different SPRTN variants with SUMO-1 in U2OS cells by confocal microscopy (Figure 3B) and employed Pearson’s correlation coefficient to quantify the co-localisation of the two signals. While a modest average correlation of 0.1 was detected between the wild-type SPRTN and SUMO-1, a significant increase to 0.4 was observed in cells expressing the protease-inactive variant E112A, further confirming a trapping effect (Figure 3B). Similar to E112A, the RJALS patient SPRTN Y117C variant, which is also proteolytically defective, co-localized with SUMO-1 foci (0.4). No correlation was observed in cells expressing the RJALS patient SPRTN variant lacking the C-terminal half (1-241), suggesting a possible involvement of the SPRTN C-terminal part in mediating interactions with SUMOylated proteins. We noticed an increase in the triple co-localisation of SPRTN-SUMO-EdU in cells expressing the inactive SPRTN variant E112A (Figure S4C, quantified in Figure S4D), suggesting that, at least in part, SPRTN processes SUMO-1 conjugates at sites of active replication.

**Figure 3.**
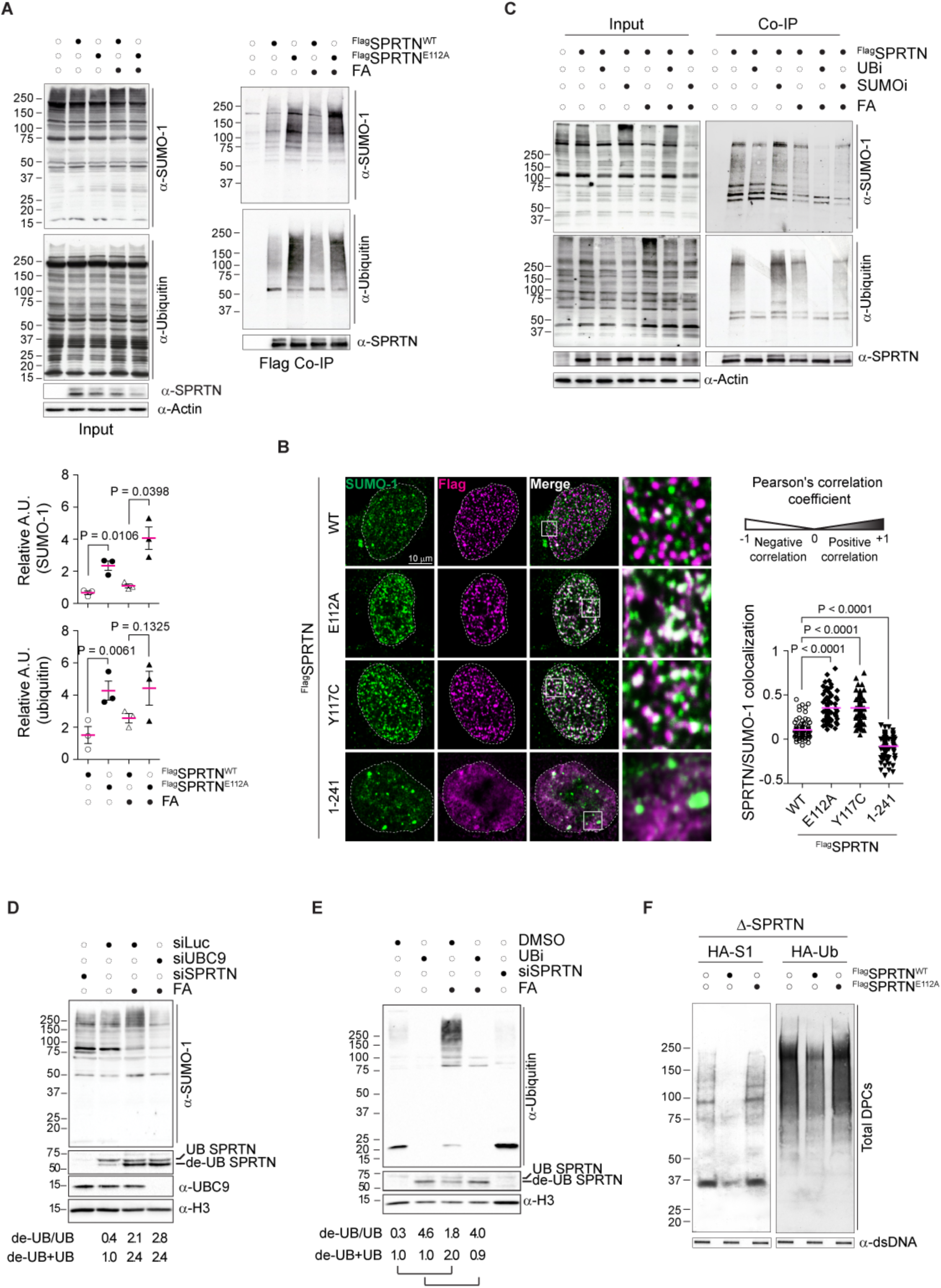
**SPRTN binds and processes SUMOylated substrates *in vivo*** (A) The catalytically inactive SPRTN^E112A^ traps SUMOylated and ubiquitylated proteins. Flag-SPRTN^wt^ or Flag-SPRTN^E112A^ were immuno-precipitated from HEK293 native total cell3 extracts. Where indicated, cells had been treated with 1 mM FA for 1 hour. Plots indicate changes in the arbitrary units (A.U.), relatively to untreated control. (B) SPRTN partially co-localises with SUMO-1 foci. U2OS cells overexpressing Flag-SPRTN^wt^, Flag-SPRTN^E112A^, Flag-SPRTN^Y117C^ and the truncated version lacking the C-terminal half (1-241) were labeled with EdU (30 minutes), pre-extracted, fixed and immunostained with the indicated antibodies. The plot indicates changes in Pearson’s correlation coefficient calculated automatically using ZEISS ZEN Microscope Software (50 nuclei). (C) SPRTN interacts with SUMOylated species in an ubiquitin-dependent manner. Flag-SPRTN^wt^ was immuno-precipitated from HEK293 native extracts after FA treatment (1 mM FA for 1 hour) in presence of the ubiquitination inhibitor MLN7243 (5 μM) or SUMOylation inhibitor 2-D08 (25 μM). (D) FA-dependent SPRTN recruitment to chromatin is not affected by blockage of the SUMOylation pathway. siLuc- or siUBC9-silenced HEK293 cells were treated, where specified, with 1 mM FA for 2 hours. Chromatin was isolated and recruitment of endogenous SPRTN was analysed by western blot. Quantification of the “ubiquitin switch” (de-UB/UB) and total amount (de-UB+UB) is reported. Representative of at least 3 independent replicates. (E) FA-dependent SPRTN recruitment to chromatin is hampered by ubiquitination inhibition. HEK293 cells were treated, wherever specified, with 1 mM FA for 2 hours in presence of DMSO or the ubiquitination inhibitor MLN7243 (5 μM). Chromatin was isolated and recruitment of endogenous SPRTN was analysed by western blot. Quantification of the “ubiquitin switch” (de-UB/UB) and relative increase in total amount (de-UB+UB) (compare DMSO vs DMSO FA; MLN7243 vs MLN7243 FA) is reported. Representative of at least 3 independent replicates. (F) SPRTN cleaves modified DPCs in cells. DPCs were isolated were isolated by RADAR from ΔSPRTN HeLa cells overexpressing Flag-SPRTN^wt^ or Flag-SPRTN^E112A^. Ectopic HA-SUMO1 (HA-S1) or HA-Ubiquitin were co-expressed together with either one of the SPRTN variants. PTMs were analysed by western blot.

Recently, it was shown that SPRTN fails to form nuclear foci following FA in the presence of the ubiquitination inhibitor MLN7243 (Borgermann et al., 2019). We recapitulated these results and found that MLN7243-treated cells (UBi) showed a reduced correlation between SPRTN and SUMO-1 foci (Figure S4E). We thus hypothesized that ubiquitin is required for SPRTN-SUMO-1 interaction. Indeed, chemical inhibition of ubiquitination prevented co-immunoprecipitation of SPRTN with SUMOylated proteins, especially after FA treatment (Figure 3C). To further explore this observation, we performed chromatin fractionations to analyse SPRTN recruitment to chromatin after FA (Stingele et al., 2016) in cells depleted of the SUMO conjugating enzyme Ubc9 or treated with MLN7243 (Figure 3D and 3E). The “SPRTN ubiquitin switch”, a phenomenon where the mono-ubiquitinated form of SPRTN is lost after FA treatment (Stingele et al., 2016), and SPRTN recruitment to chromatin were not affected by UBC9 silencing (Figure 3D, S4F). Inhibition of ubiquitin alone increased SPRTN recruitment to chromatin and caused accumulation of its de-ubiquitylated form (lower band). However, inhibition of ubiquitin in the presence of FA did not cause further SPRTN chromatin recruitment nor did it enhance the ubiquitin switch (Figure 3E, S4G). These results suggest that ubiquitin, but not SUMO, is important for SPRTN recruitment to chromatin after FA treatment. We conclude that SPRTN interacts with SUMOylated proteins in an ubiquitin-dependent manner and that ubiquitin is important for SPRTN recruitment to FA-induced chromatin lesions.

We then tested if SPRTN processes modified DPCs in cells. To this end we transiently expressed either the wild-type or the E112A SPRTN variant in ΔSPRTN cells (Figure 3F). Expression of the wild-type SPRTN reverted accumulation of SUMOylated and, in part, ubiquitinated DPCs; in contrast, the inactive mutant (E112A) failed to do so (Figure 3F). Similarly, we observed that SPRTN wild-type processed modified Topo-1 following recovery from CPT treatment, whereas the modified forms persisted in cells expressing the E112A variant (Figure S4H). We further explored SPRTN requirements for Topo-1 processing by performing *in vitro* experiments. SPRTN was not able to cleave Topo-1 after *in vitro* SUMOylation and heat-denaturation (Figure S4I). However, when purified SPRTN was incubated with Topo-1 isolated from HEK293 cells under denaturing conditions, Topo-1 and its modified forms were cleaved (Figure S4J). SPRTN protease was not able to cleave SUMO2/3 chains (Figure S4K); thereby we exclude that SPRTN is a SUMO peptidase. Considering recent reports showing that Topo-1 is modified by both ubiquitin and SUMO-1 after CPT treatment (Sun et al., 2019), our results suggest that SPRTN cleaves SUMOylated DPCs that also contain ubiquitin moieties.

### SUMO and ubiquitin are required for Replication-coupled DPC proteolysis

The above results imply that SUMO-1 and ubiquitin are both required for SPRTN-dependent replication-coupled DPC proteolysis. To test this directly, we monitored DPC removal in synchronous S-phase cells treated with SUMOylation (2-D08) or ubiquitination (MLN7243) inhibitors. Similar to SPRTN-deficient cells (Figure S3A), cells exposed to either ubiquitination or SUMOylation inhibitors failed to repair DPCs during S-phase progression (Figure 4A). Analysis of DPC modifications showed that, in SUMOylation-deficient cells, both SUMO and ubiquitin signals persisted during S-phase progression and were only partially removed from the DPCs (Figure 4B). In MLN7243-treated cells, SUMO-1-DPCs persisted, further supporting the need for ubiquitination in processing SUMOylated DPCs. In the soluble fraction, inhibition of SUMOylation increased the ubiquitin signal and vice versa (Figure S5A), again hinting a possible competition between SUMO and ubiquitin. Next, we exposed cells to low doses of CPT and monitored Topo-1cc repair. Similar to SPRTN deficient cells (ΔSPRTN), inhibition of either ubiquitination or SUMOylation led to accumulation of TOPO-1ccs (Figure 4C). No additive effect was detected upon double inhibition, suggesting that ubiquitin and SUMO work in the same pathway (Figure 4C, right panel). Likewise, hampering SUMOylation by UBC9 depletion also contributed to Topo-1cc accumulation (Figure S5B). No additive effect was detected upon co-depletion with SPRTN (Figure S5B, bottom panel), suggesting that both SUMO and SPRTN work in the same pathway. Altogether, these results show the requirement for both ubiquitin and SUMO in replication-coupled DPC proteolysis repair.

**Figure 4.**
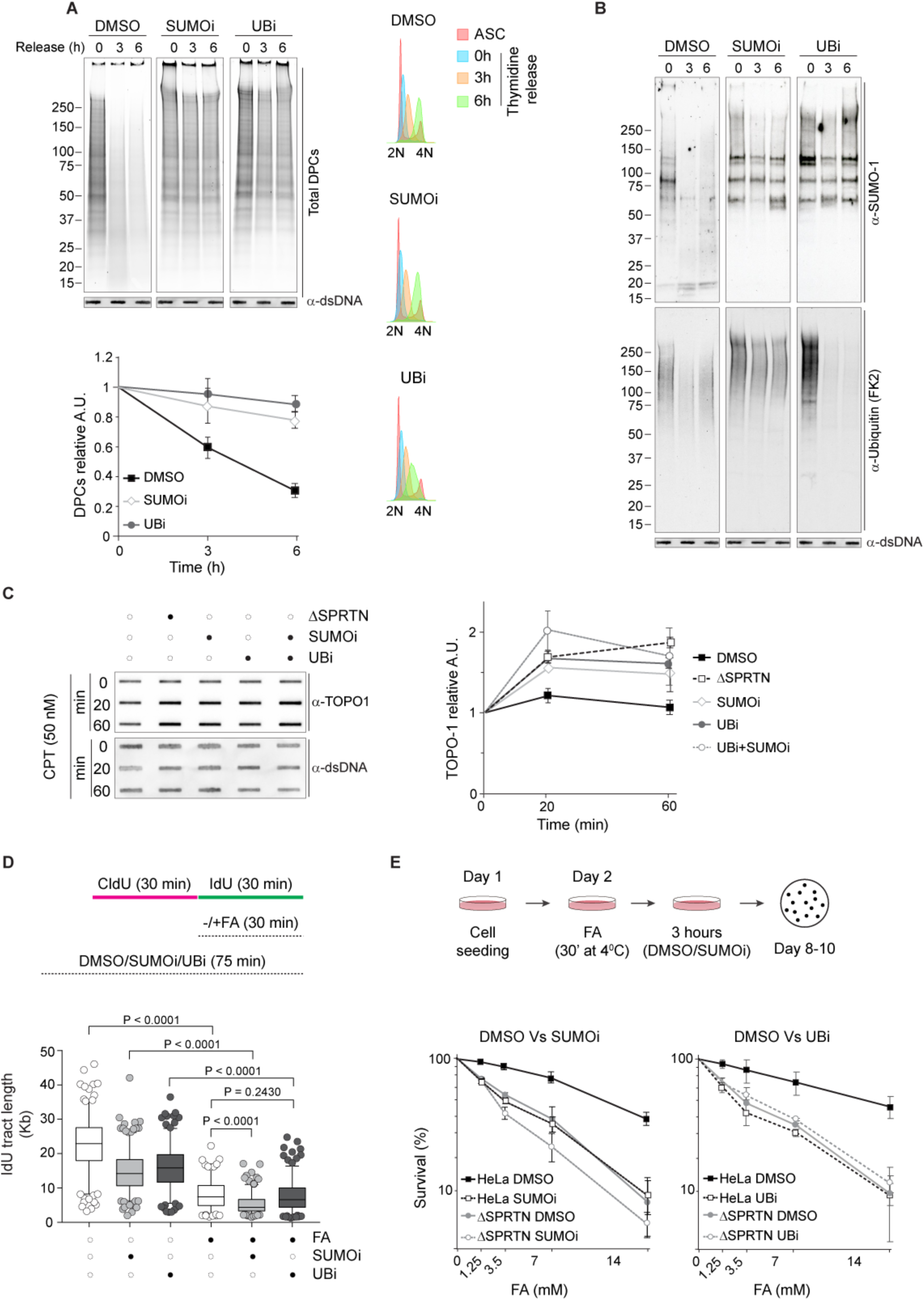
**SUMO and ubiquitin are required for Replication-coupled DPC proteolysis** (A) SUMOylation and ubiquitination inhibition block DPC removal during S-phase progression. HeLa cells were synchronized in G1/S with double thymidine block and released in the presence of DMSO, 25 μM 2-D08 (SUMOi) or 5 μM MLN7243 (UBi). Total DPCs were isolated by RADAR and detected by Flamingo protein gel staining. Slot blot with anti-dsDNA was used as a loading control. Right panel shows cell cycle distribution by FACS analysis of the DNA content (propidium iodide). (B) SUMOylated DPCs persist upon ubiquitination inhibition. DPCs isolated in (A) were analysed by western blot for SUMO-1 and ubiquitin. (C) SUMOylation and ubiquitination inhibition hamper Topo-1cc resolution. HeLa cells were treated with 50 nM CPT for the indicated times in presence of the ubiquitination inhibitor MLN7243 (5 μM) or SUMOylation inhibitor 2-D08 (25 μM). Total DPCs were isolated by RADAR and Topo-1ccs were analysed by slot blot. dsDNA was used as a loading control. Graph (right panel) shows the mean ± SEM of the relative signal from 2 independent experiments. (D) SUMOylation and ubiquitination inhibition reduce DNA replication speed. Box and whiskers plot for DNA combing analysis. HEK293 cells were allowed to incorporate CldU for 30 min and IdU for additional 30 min in the presence of DMSO, 50 μM 2-D08 (SUMOi) or 5 μM MLN7243 (UBi). Where indicated, 450 μM FA was added for the duration of IdU incubation. The length of IdU tracts was measured with FiberVision Software and statistical significance calculated using unpaired t-test (Mann-Whitney). (E) SUMOylation and ubiquitination inhibition sensitize cells to FA. Schematic of the survival assay protocol (upper panel). Parental or ΔSPRTN HeLa cells were exposed to the indicated concentrations of FA for 30 min at 4°C and let recover for 4 hours in the presence of DMSO, 25 μM 2-D08 (SUMOi) or 5 μM MLN7243 (UBi). Colonies were allowed to grow for 8-10 days before fixation and counting. Graphic representation of the survival fraction from 2 independent experiments.

### Functional SUMOylation and ubiquitination responses promote genomic stability

Considering that SUMO and ubiquitin are both required for replication-coupled DPC proteolysis, we predicted that inhibition of both these pathways would exacerbate DNA replication defects after DPC induction. To test this idea, we monitored the fitness of replication fork progression by using a DNA combing assay (Figure 4D and S5C). As previously reported, DNA track length was reduced by FA treatment (Halder et al., 2019; Mórocz et al., 2017; Vaz et al., 2016). We also observed a reduction when cells were exposed to SUMOylation or ubiquitination inhibitors. When cells were exposed to FA in the presence of either one of the two inhibitors, DNA track length was further reduced, suggesting the need for both SUMO and ubiquitin signals in the progression of the DNA replication fork over DPCs (Figure 4D and S5C).

To explore the role of SUMO and ubiquitin in preventing DPC-induced toxicity, we assessed cellular sensitivity to FA and CPT. Similar to SPRTN deficient cells (ΔSPRTN), SUMOylation inhibition sensitised wild-type cells to both DPC-inducing agents (Figure 4E and S5D). Ubiquitination inhibition produced similar results (Figure 4E). The sensitivity was not further increased in ΔSPRTN cells. Altogether, this suggests that SUMOylation, ubiquitination and SPRTN work in the same pathway for DPC repair.

### SUMO suppresses excessive ubiquitin signal at DPC-induced DNA damage sites

While investigating the ubiquitin and SUMO signals at DNA damaged sites, we noticed that these two PTMs appeared to be mutually exclusive in SPRTN-deficient cells (Figure 2B and 2C). In light of recent reports showing a proteasome-dependent DPC repair mechanism in *Xenopus* egg extracts (Larsen et al., 2019; Sparks et al., 2019), we set out to test whether the 26S proteasome might function as the proteolytic alternative to SPRTN. Total DPC removal kinetics were not delayed by the proteasome inhibitor MG132 following recovery from a short pulse with FA (Figure S6A), although we did observe persistence of ubiquitin signal on DPCs at the 3-hour recovery point (Figure S6B). Increase of ubiquitin signal in high molecular weight regions of soluble proteins in MG132 treated cells in 60 and 180 mins of recover time indicates that proteasome inhibition worked (Figure S6C). Likewise, analysis of Topo-1ccs following treatment with low doses of CPT showed that the proteasome inhibition did not interfere with Topo-1cc removal (Figure S6D). We therefore conclude that the proteasome has a negligible role in DPC removal under our experimental conditions.

We decided to analyse further the interplay between SUMO and ubiquitin at DPC-induced damage sites by blocking SUMOylation. SUMOylation-deficient cells showed a 3-fold increase in ubiquitin foci following a 3 hour-recovery from short FA pulse (Figure 5A). Co-localisation of ubiquitin with γH2AX indicates that ubiquitin foci formed at FA-induced damage sites (Figure 5B). Similarly, SUMOylation inhibition led to a robust increase in the ubiquitin signal at DNA damage sites following recovery from UV laser micro-irradiation (Figure 5C). These results suggest that SUMOylation suppresses excessive ubiquitination at DPC-induced DNA damage sites.

**Figure 5.**
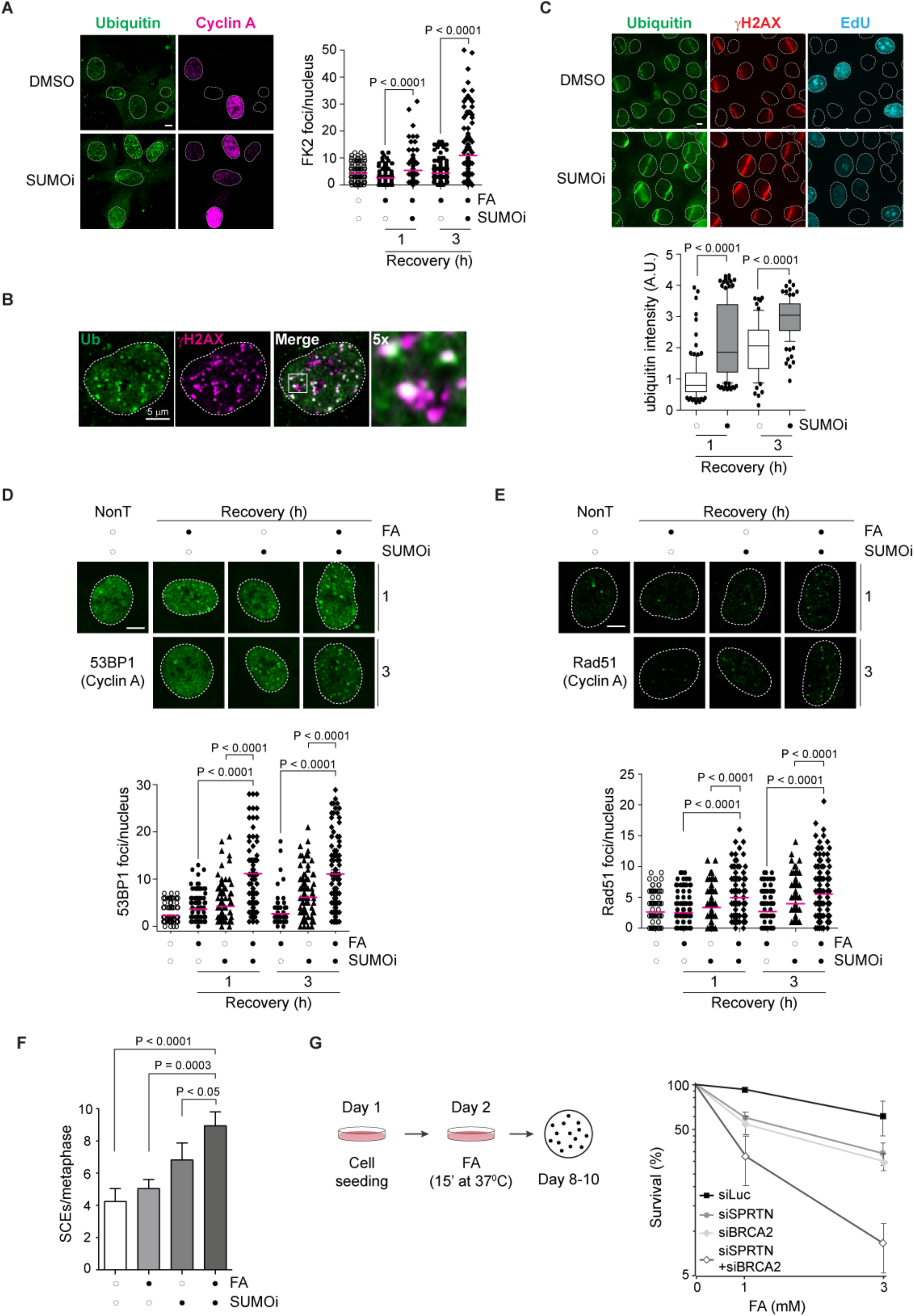
**SUMO suppresses excessive ubiquitin signal at DPC-induced DNA damage sites** (A) SUMOylation inhibition induces ubiquitin foci following FA treatment. RPE-1 cells were treated with 1 mM FA for 20 minutes at 4°C and let recover for the indicated time-points in the presence of DMSO or 25 μM 2-D08. After treatment, cells were fixed and immunostained with the indicated antibodies. Left panel: representative picture at 3 hours after treatment. Right panel: graphic representation of the number of foci per nucleus (100 nuclei) counted with ImageJ. Statistical significance was calculated using unpaired t test. (B) Ubiquitin foci form at DNA damage sites (γH2AX). Confocal microscopy of RPE-1 cells treated as described above. 3 hours after treatment cells were fixed and immunostained with the indicated antibodies. (C) SUMOylation inhibition increases total ubiquitin signal at DNA damage sites. UV laser microirradiation was performed in HeLa cells as described in Figure 2C. Following DNA damage, cells were allowed to recover in the presence of DMSO or 25 μM 2-D08 (SUMOi) for the indicated times. Cells were then pre-extracted, fixed and immunostained with the indicated antibodies. Signal at DNA damage sites was quantified using ImageJ and statistical significance calculated using unpaired t-test. (D) SUMOylation inhibition increases 53BP1 foci formation. RPE-1 cells were treated with 1 mM FA for 20 minutes at 4°C and allowed to recover for the indicated times in presence of DMSO or 25 μM 2-D08 (SUMOi). After treatment, cells were fixed and immunostained with the indicated antibodies. Upper panel: representative images. Lower panel, graphic representation of the number of 53BP1 foci per nucleus in cyclin A-positive cells (100 nuclei) counted using ImageJ. Statistical significance was calculated using unpaired t-test. (E) SUMOylation inhibition increases Rad51 foci formation. RPE-1 cells were treated with 1 mM FA for 20 minutes at 4°C and allowed to recover for the indicated times in presence of DMSO or 25 μM 2-D08 (SUMOi). After treatment, cells were fixed and immunostained with the indicated antibodies. Upper panel: representative images. Lower panel, graphic representation of the number of Rad51 foci per nucleus in cyclin A-positive cells (100 nuclei) counted using ImageJ. Statistical significance was calculated using unpaired t-test. (F) SUMOylation suppresses sister chromatid exchange events (SCEs). HeLa cells were grown for 48 hours in presence of BrdU, exposed to 450 μM FA for 10 minutes at 37°C with 10 μM 2-D08 (SUMOi) or DMSO and allowed to recover for 16 hours in the presence of colcemid with DMSO or 10 μM 2-D08. Metaphase spreads were stained and SCE events counted from at least 30 nuclei. (G) Co-depletion of SPRTN and BRCA2 hyper-sensitizes cells to FA. Schematic of the survival assay protocol (left panel). Depleted cells were exposed to the indicated concentrations of FA for 15 min at 37°C. Colonies were allowed to grow for 8-10 days before fixation and counting. Graphic representation of the survival fraction from 2 independent experiments.

In yeast, SUMO supresses recombinogenic events (Branzei et al., 2006). As HR is also active in S-phase, we wondered: (i) whether the late ubiquitination signal is associated with a recombinogenic pathway, and (ii) if DPC persistence in SUMOylation-defective cells switches the DPC repair pathway from SPRTN-dependent proteolysis to HR. Indeed, SUMOi-treated RPE-1 cells showed an increase in phospho-RPA, phospho-CHK2 and phospho-KAP1, the well-defined markers of resected ssDNA and DNA double-strand breaks (DSBs), following treatment with FA or CPT (Figure S7A). On laser stripes, ΔSPRTN cells accumulated more SUMO-1 (Figure 2C) and less RPA 1 hour after UV laser micro-irradiation (Figure S7B). Strikingly, at the 9-hour recovery point, RPA signal increased in ΔSPRTN cells, correlating with a lower SUMO-1 signal and a high ubiquitin signal (Figure 2C and S7B).

To confirm that FA treatment causes DSBs in SUMOylation-deficient cells due to defective SPRTN-dependent proteolysis, we monitored Rad51 and 53BP1 foci, two well-recognised markers for DSB formation, by immunofluorescence microscopy. Cells recovering from FA treatment in the presence of SUMOylation inhibitor showed a 4-fold increase in the average number of 53BP1 foci (Figure 5D) and a 2-fold increase in the average number of Rad51 foci per nucleus (Figure 5E). We observed an increase in sister chromatid exchange (SCE) events when cells were treated with FA and allowed to recover in the presence of the SUMO inhibitor (Figure 5F and S7C). Collectively, this set of data supports a role for SUMO in DPC repair by promoting SPRTN-dependent proteolysis and suppressing an ubiquitin-dependent, recombinogenic back-up pathway.

Two additional pieces of evidence confirmed that HR is a back-up for DPC repair in SPRTN-deficient cells. Firstly, depletion of the HR factor BRCA2 further sensitised SPRTN-depleted cells to FA (Figure 5G). Secondly, SPRTN depletion by siRNA in BRCA2-deficient cells inhibited growth and caused cell death, suggesting synthetic lethality (Figure S7D).

Taken together, our results show that the SPRTN-SUMO axis protects cells from DPC-induced DSBs. SUMOylation promotes SPRTN-dependent DPC proteolysis in order to counteract excessive ubiquitination and activation of an HR-dependent recombinogenic repair pathway.

## DISCUSSION

Our results reveal a crosstalk between SUMO-1 and ubiquitin in the replication-coupled DPC repair pathway, providing new insights into the regulatory signaling mechanisms associated with DPC proteolysis. We propose that SUMO-1 favors SPRTN-dependent proteolysis over a recombinogenic pathway. We demonstrated that clearance of DPCs during DNA replication is a SUMO/ubiquitin/SPRTN-dependent process essential for DNA replication fork progression and genome stability. Our conclusion is based on the following observations: (1) SUMO-1 accumulates in nuclear foci and on DPCs in SPRTN-deficient cells; (2) SPRTN interacts with SUMOylated proteins in a ubiquitin-dependent manner; (3) SUMO and ubiquitin promote DPC clearance during DNA replication allowing unperturbed DNA replication fork progression and preventing DPC-induced cytotoxicity; (4) SUMO suppresses excessive ubiquitination at sites of DNA damage and on DPCs; (5) SUMO prevents DPC-induced recombinogenesis and cell lethality after exposure to DPC inducing agents.

SPRTN and its yeast ortholog Wss1 show pleiotropic activity on DNA-binding proteins *in vitro* (Mórocz et al., 2017; Stingele et al., 2014, 2016; Vaz et al., 2016), while it is predicted that their activities are regulated *in vivo* to avoid uncontrolled cleavage of nuclear proteins (Fielden et al., 2018; Stingele and Jentsch, 2015). Strong transient overexpression of SPRTN has indeed been found to be toxic (Lessel et al., 2014). Considering that SPRTN travels with the replication fork (Maskey et al., 2017; Vaz et al., 2016), replisome components are at particularly high risk of potentially undesired cleavage. Therefore, the labelling of potential SPRTN substrates with PTMs arises as a plausible regulatory mechanism to prevent uncontrolled proteolysis.

It is generally accepted that SPRTN and Wss1 activities are regulated by ubiquitin and SUMO, respectively (Stingele et al., 2015). SPRTN has a UBZ domain for binding to ubiquitin and Wss1 has several SUMO-interacting motifs. However, by analysing SUMO and ubiquitin on the DPCs of SPRTN-deficient cells, we found that SPRTN prevents accumulation of SUMO-1 but not ubiquitin (Figure 2A). In fact, ubiquitination appears to be a transient modification even in SPRTN-deficient cells (Figure 2A, lower panel), hinting at a signaling event whose nature is elusive at the moment. Our data suggest that ubiquitin is at least required for the interaction between SPRTN and SUMO-1 substrates. Ubiquitin and SPRTN UBZ are also required for SPRTN recruitment to DNA damage sites (Centore et al., 2012; Davis et al., 2012; Ghosal et al., 2012; Machida et al., 2012; Mosbech et al., 2012) and for SPRTN-dependent proteolysis in *Xenopus* egg extract (Larsen et al., 2019).

It seems plausible that both SUMO and ubiquitin co-exist on the same substrate. In fact, our *in vitro* cleavage experiments on Topo-1 indicate that SPRTN efficiently cleaves Topo-1 only when isolated from CPT-treated cells, a condition that induces co-modification by SUMO and ubiquitin (Sun et al., 2019). Such crosstalk between ubiquitin and SUMO could signal SPRTN ubiquitin-dependent proteolysis of SUMOylated DPCs, thus preventing unscheduled cleavage of proteins modified by only one of the two PTMs.

Recent studies in a *Xenopus* cell-free system implicated the proteasome in the degradation of ubiquitylated DPCs (Larsen et al., 2019). Nonetheless, in our system proteasome inhibition does not affect the repair of either total DPCs (Figure S6A) or Topo-1ccs (Figure S6D), largely excluding this proteolytic pathway from replication-coupled repair. These differences may be due to the excess of proteins in *Xenopus* egg extracts compared to cell-based systems and/or the lower complexity of the plasmid-DPC investigated in *Xenopus* cell-free extracts compared to the high complexity of chromatin in the cell. Other reports also implicated the proteasome in DPC – Topo1/2-cc – proteolysis (Desai et al., 1997; Lin et al., 2008; Mao et al., 2001). However, caution should be taken in interpreting these results, since long treatments with proteasome inhibitors can deplete the nuclear pool of ubiquitin (Mimnaugh et al., 1997), and therefore DPC persistence could be an indirect effect. Alternatively, the proteasome could come into play when DPC formation extends beyond the SPRTN-dependent repair capacity. In fact, proteasome inhibition causes TOPO-1cc accumulation only when cells are exposed to high doses of CPT (20μM or higher) (Desai et al., 1997; Interthal and Champoux, 2011; Lin et al., 2008; Mao et al., 2001; Sordet et al., 2009) rather than the low and clinically relevant doses (25nM) used in our experimental set-up.

In addition to the initial transient signal, we found that SPRTN depletion and SUMOylation inhibition lead to a late ubiquitin accumulation at DNA damage sites (Figure 2C and 5C). We noticed that the accumulation of SUMO-1 and ubiquitin are mutually exclusive (Figure 2C). Furthermore, the increase in ubiquitin signal mirrors increased accumulation of replication protein A (RP-A) and Rad51 together with increased number of 53BP1 foci, suggesting the formation of DSBs and activation of the HR pathway (Figure 5D, 5E and S7B). Based on these findings, we propose that SPRTN/SUMO1-dependent proteolysis is the preferential pathway for DPC repair. Recruitment and regulation of several HR proteins, including RAP80-Abraxas, BRCA1 and Rad51, heavily relies on ubiquitin (Schwertman et al., 2016). In fact, we did observe the occurrence of a strong ubiquitin signal when SUMOylation is inhibited. We conclude that ubiquitin-dependent HR is a back-up pathway for DPC repair during S-phase but at the cost of chromosomal rearrangements.

How SUMOylation of DPCs/DNA lesions suppresses ubiquitination signal in order to foster SPRTN-dependent pathway is not clear. However, our results are in line with previous observations made in yeast, showing that SUMOylation counteracts recombinogenic events at damaged replication forks (Branzei et al., 2006), and reinforce the need for the SUMO pathway during DNA replication to cope with DPCs. The wide range of possibilities for the interplay between SUMO and ubiquitin in DPC repair will require an extensive analysis that goes beyond the scope of this paper. Nonetheless, our data establish a role for this crosstalk in DPC proteolysis repair, cell survival and prevention of genomic rearrangements.

## Acknowledgements

This work was supported by the Medical Research Council-UK (MC_EX_MR/K022830/1) and Oxford Cancer Research Centre to K.R. A.R. is supported by an EMBO long-term fellowship (ALTF 1109-2017). The authors thank members of the Ramadan lab for discussions.

## Author contributions

B.V. performed DPC isolations, microscopic analysis of nuclear foci, clonogenic assays and SPRTN immunoprecipitation experiments; A.R. performed chromatin fractionations and DNA combing; M.P., Topo-1 *in vitro* cleavage experiments. A.R. and G.R.-B., SCEs experiments; S.K., laser stripe experiment in Figure 2C. A.N.S. purified recombinant SPRTN. B.V., A.R. and K.R wrote the manuscript. All authors discussed and analysed the data. K.R. conceived and supervised the project.

---

**Supplementary figure 1. Related to Figure 1.**

(A) FA treatment promotes ubiquitination and SUMOylation on soluble proteins. HeLa cells were treated and processed as in Figure 1A. Protein soluble fractions were analysed by western blot for the indicated post-translational modifications (PTMs).
(B) Total DPCs do not increase with longer FA treatments. HeLa cells were exposed to 1.35 mM FA at 37°C for the indicated times. Total DPCs were isolated by RADAR and analysed by SDS-PAGE followed by Flamingo protein gel staining. Slot blot with anti-dsDNA was used as a loading control.
(C) FA does not activate the Fanconi anemia pathway. HEK293 cells were treated with 1 mM FA for 20 minutes at 4°C and recovered for the indicated times, or treated continuously for 2 hours. Alternatively, cells were treated with 1 μM Mitomycin C (MMC) or 5 μM cisplatin (Cis) for 24 hours.
(D) Interstrand crosslinking agents do not induce DPCs. HeLa cells were exposed to 1 μM Mitomycin C (MMC) or 5 μM cisplatin (Cis) for 2 hours. Total DPCs were isolated by RADAR and analysed by SDS-PAGE followed by Flamingo protein gel staining. Slot blot with anti-dsDNA was used as a loading control.
(E) CPT treatment promotes formation of Topo-1 crosslinks. HeLa cells were treated with increasing concentrations of CPT for 15 minutes at 37°C. DPCs were isolated as described in Figure 1A and analysed by either Flamingo protein gel staining (left panel) or by immuno-blot against Topo-1 (right panel). Slot blot with anti-dsDNA was used as a loading control.
(F) CPT treatment promotes ubiquitination and SUMOylation (SUMO-2/3) in soluble and DPC fractions. HeLa cells were treated as in (E). Soluble and DPC proteins were analysed by immuno-blot with respective antibodies against different PTMs.
(G) FA treatment increases ubiquitin pan-nuclear staining. Quantification of the ubiquitin signal intensity using ImageJ (from Figure 1C).
(H) FA treatment induces DPCs mostly in Asynchronous (ASC) RPE-1 cells. RPE-1 cells were treated with 1.35 mM FA for 10 minutes at 37°C. Total DPCs were isolated by RADAR and detected with Flamingo protein gel staining. Samples were immuno-blotted with the indicated antibodies. Slot blot with anti-dsDNA was used as a loading control. Cell cycle distribution by FACS (far left panel) analysis of the DNA content (propidium iodide) confirms cell cycle arrest of G0 cells.
(I) DPC modification is increased in the presence of HU. HeLa cells were exposed to HU (1 mM) for 1 hour when indicated. FA was added (1.35 mM) and cells were incubated for 10 minutes at 37°C. Total DPCs were isolated by RADAR and PTMs on DPCs analysed by immuno-blot with the indicated antibodies.

**Supplementary figure 2. Related to Figure 2.**

(A) SPRTN-deficient cells show a slower DPC repair kinetics. Parental and ΔSPRTN HeLa cells were exposed to 1.35 mM FA for 10 minutes at 37°C and allowed to recover for the indicated time-points (as in Figure 2A). Total DPCs were isolated by RADAR and detected with Flamingo protein gel staining. Representative picture (upper panel) and graphic (lower panel) of the mean ± SEM of relative DPC signal from 3 independent experiments measured with ImageJ. Slot blot with anti-dsDNA was used as loading control.
(B) SUMOylation wave persists in SPRTN-depleted cells. Cells were treated and processed as in Figure 2A. Total DPCs were analysed by western blot with anti-SUMO-2/3. Graph shows the mean ± SEM of relative signal from 3 independent experiments.
(C) SPRTN deficiency (ΔSPRTN cells) does not affect cell cycle distribution. Parental and ΔSPRTN HeLa cells were treated as in Figure 2A. Cell cycle distribution of the different phases was determined by FACS analysis of the DNA content and EdU labeling.
(D) SPRTN deficiency (ΔSPRTN cells) does not affect significantly DNA replication. Parental and ΔSPRTN HeLa cells were treated as in Figure 2A and EdU mean intensity determined by FACS analysis.
(E) Ubiquitin and SUMO conjugates are resolved in the soluble fraction of both parental and ΔSPRTN HeLa cells following FA treatment. Soluble proteins were isolated as described in Figure 1A from cells treated as in Figure 2A. Samples were analysed by immuno-blot with the indicated antibodies.

**Supplementary figure 3. Related to Figure 2.**

(A) ΔSPRTN cells accumulate DPC during S-phase. HeLa parental or ΔSPRTN cells were synchronized at G1/S phase of cell cycle and released. Total DPCs were isolated by RADAR and detected with Flamingo protein gel staining. Lower panel indicates cell cycle progression by FACS.
(B) ΔSPRTN cells are defective in removal of SUMO-DPCs during S-phase. DPCs from Figure S3A were analysed by immunoblot with the indicated antibodies.
(C) SPRTN-deficient cells accumulate SUMO-1, partially at DNA damage sites. SPRTN was depleted by siRNA in U2OS cells for 72 hours. Cells were labeled with EdU, pre-extracted, fixed and immuno-stained with the indicated antibodies.
(D) RJALS patient cells accumulate SUMOylated DPCs. Normal and patient lymphoblasts were treated as in Figure 2A and recovered for the indicated time. Chromatin was isolated and SUMO-1 signal was analysed by western blot. Right panel: quantification of SUMO-1 signal in reference to time-point 0.

**Supplementary figure 4. Related to Figure 3.**

(A) SPRTN immunoprecipitates contain SUMO-2/3 moieties. HEK293 cells expressing Flag-SPRTN^wt^ or Flag-SPRTN^E112A^ were treated and processed as in Figure 3A. Immunoprecipitates were analysed by immunoblot with anti-SUMO2/3 antibodies.
(B) SPRTN interacts with SUMOylated and ubiquitylated proteins. SUMO- and ubiquitin-conjugated proteins were released from SPRTN immunoprecipitates after incubation in highly stringent conditions (1 M NaCl).
(C) The inactive SPRTN protease (E112A) accumulates at SUMO-1- and EdU-positive sites. U2OS overexpressing either the Flag-SPRTN^wt^ or the Flag-SPRTN^E112A^ were labelled with EdU (10 minutes), and processed as in Figure 3B.
(D) Pearson’s correlation coefficient changes for Figure S4C. Correlation coefficients were calculated as in Figure 3B (50 nuclei).
(E) MLN7243 treatment reduces co-localisation between SPRTN and SUMO-1 foci. U2OS cells expressing Flag-SPRTN^wt^ were treated with 1 mM FA in the presence or absence of 5 µM MLN7243 (UBi). 1 hour after treatment cells were pre-extrated, fixed and immunostained with the indicated antibodies. Right panel shows changes in Pearson’s correlation coefficient.
(F) FA-dependent SPRTN “ubiquitin switch” is not affected by blockage of the SUMOylation pathway. Total cell extracts from experiment in Figure 3D. Asterisk indicates an unspecific band.
(G) Ubiquitination inhibitor hampers the FA-dependent SPRTN “ubiquitin switch”. Total cell extracts from experiment in Figure 3E. Asterisk indicates an unspecific band.
(H) Cells expressing SPRTN^E112A^ accumulate modified Topo-1 after CPT treatment. ΔSPRTN cells expressing either Flag-SPRTN^wt^ or Flag-SPRTN^E112A^ were treated for 1 hour with CPT (1 μM) and allowed to recover for 1 hour. Chromatin was isolated with the indicated conditions and analysed by western blot.
(I) SPRTN does not cleave SUMOylated Topo1 *in vitro*. Recombinant Topo-1 was SUMOylated *in vitro*, denatured and incubated with *E.coli*-purified SPRTN for 16 hours at 37°C.
(J) SPRTN cleaves modified TOPO-1 *in vitro*. YFP-TOPO1 was immunopurified under denaturing conditions from HEK293 after CPT treatment (10 μM CPT for 1 hour). GFP beads, containing YFP-TOPO1, were incubated with either *E.coli*-purified SPRTN^wt^ or SPRTN^E112A^ for 16 hours at 37°C and reactions were analysed by western blot. Arrowheads indicate cleavage products; asterisks indicate cross-reactive bands.
(K) SPRTN is not a SUMO peptidase. SUMO-2/3 chains were incubated with either *E.coli*-purified SPRTN^wt^ or SPRTN^E112A^ and reactions were analysed by western blot.

**Supplementary figure 5. Related to Figure 4.**

(A) Analysis of the SUMOylated and ubiquitylated proteins in the soluble fraction of the cells analysed in Figure 4A.
(B) Co-depletion of SPRTN and UBC9 does not have an additive effect in Top1cc removal. siSPRTN and siUBC9 HeLa cells were treated with 50 nM CPT for the indicated times. Total DPCs were isolated by RADAR and Topo-1ccs were analysed by slot blot. dsDNA was used as a loading control. Graph shows the mean ± SEM of the relative signal from 3 independent experiments.
(C) SUMOylation and ubiquitination inhibition reduce DNA replication speed. Representative images from experiment in Figure 4D.
(D) SUMOylation inhibition sensitizes cells to CPT. Parental and ΔSPRTN HeLa cells were exposed to the indicated concentrations of CPT for 4 hours in presence of DMSO or 25 μM 2-D08. Colonies were allowed to grow for 8-10 days before fixation and counting. Graphic representation of the survival fraction from 3 independent experiments.

**Supplementary figure 6. Related to Figure 5**.

(A) Proteasome inhibition does not impair DPC removal. HeLa cells were treated with 1.35 mM FA for 10 minutes at 37°C and allowed to recover for the indicated times in presence of DMSO or 10 μM MG132. Total DPCs were isolated by RADAR and detected with Flamingo protein gel staining.
(B) Proteasome inhibition does not affect removal of SUMO-1 DPCs. HeLa cells were treated as described in (A). DPCs were isolated by RADAR and analysed by western blot for SUMO-1 and ubiquitin modifications. Slot blot with anti-dsDNA was used as loading control.
(C) Proteasome inhibition causes accumulation of ubiquitin conjugates in the soluble fraction following FA treatment. Soluble proteins were isolated as described in Figure 1A from the samples prepared as described in (A) and analysed by western blot for the indicated PTMs.
(D) Proteasome inhibition does not affect removal of Topo-1cc. HeLa cells were treated continuously with low doses of CPT (50 nM) for the indicated times in the presence of the indicated inhibitors. DPCs were isolated by RADAR and analysed by slot blot with antibodies against Topo-1. Slot blot with anti-dsDNA was used as loading control.

**Supplementary figure 7. Related to Figure 5.**

(A) SUMOylation inhibition enhances the activation of DNA damage response markers. RPE-1 cells were treated with 1 mM FA or 50 nM CPT for 1 hour, in combination with DMSO or 25 μM 2-D08. Total cell extracts were prepared and analysed by immunoblot for the indicated proteins.
(B) RPA signal on laser stripes in ΔSPRTN cells. Parental and ΔSPRTN HeLa cells were treated as in Figure 2C, fixed and immunostained with the indicated antibodies. Bottom panel: quantification of the changes in RPA signal over time.
(C) SUMOylation suppresses sister chromatid exchange events (SCEs). Representative metaphases from experiment in Figure 5F.
(D) SPRTN depletion induces synthetic lethality in BRCA2 deficient cells. SPRTN was depleted at day 2 and day 5 by two siRNA sequences in either BRCA2-deficient (DLD1^-/-^) or complemented (DLD1^+/+^) cells. Cells were counted manually for the indicated time using trypan blue to exclude dead cells.

## STAR METHODS

### Lead contact and materials availability

Further information and material requests should be addressed to the lead contact.

### Experimental model and subject details

#### Cell culture

HEK293, HeLa, RPE-1 and U2OS cells were grown in Dulbecco’s Modified Eagle’s medium (DMEM, Sigma-Aldrich) supplemented with 10% fetal bovine serum (Sigma-Aldrich) and 100 I.U./mL penicillin - 0.1 mg/mL streptomycin (Sigma-Aldrich) at 37°C in a humidified incubator with 5% CO2, and tested for mycoplasma contamination. CRISPR partial knockout ΔSPRTN HeLa cells (Vaz et al., 2016) were maintained as above.

### Method details

#### Cellular treatments and transfections

Treatments with FA and CPT were performed as stated in the individual protocols. Transfections were performed using Lipofectamine® 3000 and expression of the genes was allowed for 24-48 hours; transfection with siRNAs was performed with Lipofectamine RNAiMAX transfection reagent and silencing was allowed for 72 hours.

### Cell cycle synchronization

HeLa cells were synchronized at G1/S of the cell cycle by double thymidine treatment, as described previously (Harper, 2005). RPE-1 cells were synchronized by contact inhibition. Cells were maintained for 48 hours at 100% confluence. HeLa cells were serum starved for 48 hours at 80-90% confluence.

### Western Blot

Standard protocols for sodium dodecyl sulfate-polyacrylamide gel electrophoresis (SDS-PAGE) and immuno-blotting were used (Henderson and Wolf, 1992). Nitrocellulose membrane (GE Healthcare) or Polyvinylidene fluoride (PVDF) (Bio-Rad) were used to transfer proteins from polyacrylamide gels depending on the antibody. Acquisition was performed with a Bio-Rad ChemiDoc XRS Plus Analyzer or X-Ray film (Scientific Laboratory Supplies). Quantification of western blot bands was performed on ImageLab™ software (Bio-Rad) or ImageJ after scanning the film.

### DPC isolation

DPCs were detected using a modified rapid approach to DNA adduct recovery (RADAR) assay (Kiianitsa and Maizels, 2013). In brief, 1.5 to 2 x 10^6^ cells were lysed in 1 ml of DPC lysis buffer, containing 6 M guanidinium isothiocyanate, 10 mM Tris–HCl (pH 6.8), 20 mM EDTA, 4% Triton X100, 1% N-Lauroylsarcosine Sodium and 1% dithiothreitol. DNA was precipitated by adding 1 ml of 100% ethanol. When indicated, the supernatant was saved for protein precipitation with 1:9 100% ethanol. The DNA pellet was washed three times in wash buffer (20 mM Tris–HCl pH 7.5, 50 mM NaCl, 1 mM EDTA, 50% ethanol). DNA was solubilized in 1 ml of 8 mM NaOH. A small aliquot of the recovered DNA was digested with 50 μg/ml proteinase K for 3 hours at 50°C. DNA concentration was determined using a Quant-iT™ PicoGreen™ dsDNA assay kit (ThermoFisher Scientific) according to manufacturer’s instructions. Normalized amounts of dsDNA containing the DPCs were digested with benzonase for 1 hour at 37°C. Proteins were precipitated by standard Trichloroacetic Acid (TCA) protocol (Link and Labaer, 2011) and resolved by SDS-PAGE gel.

### DPC detection

Total DPCs were visualized by Flamingo™ Fluorescent Protein Gel Stain (Bio-Rad) as recommended by the manufacturer after electrophoretic separation on polyacrylamide gels. Specific DPCs were detected by immuno-staining. In brief, dsDNA containing the DPCs was quantified using a Quant-iT™ PicoGreen™ dsDNA assay kit. Normalized amounts of DNA were digested with benzonase (Merck Millipore) for 1 hour at 37°C. Samples were diluted in Tris-buffered saline (TBS) and applied to a PVDF membrane using a vacuum slot-blot manifold (Bio-Rad). The membrane was then blocked in 3% BSA in TBS/T (TBS, 0.1% Tween-20) and incubated with primary antibodies followed by incubation with HRP-conjugated secondary antibodies. DPCs were visualized using a Bio-Rad ChemiDoc XRS Plus Analyzer. For slot-blot detection of dsDNA, 10-20 μg of DNA were incubated with proteinase K to digest the crosslinked proteins, diluted in Tris/Borate/EDTA (TBE) buffer and applied to nylon membrane (GE Healthcare). The membrane was blotted with an anti-dsDNA antibody and developed as before.

### UV laser microirradiation

Cells were seeded onto 10 mm No. 1 glass coverslips (VWR) 24 hours before the experiment. Twenty minutes before UV-A laser, cells on coverslips were treated with 10 μg/mL Hoechst and 10 μM EdU, and micro-irradiated using 355 nm pulsed laser connected to a Nikon TE2000 microscope. After recovery of indicated times, cells were pre-extracted on ice for 5 minutes with 25 mM HEPES (pH 7.4), 50 mM NaCl, 1 mM EDTA, 3 mM MgCl2, 300 mM sucrose, and 0.5% Triton X-100. Cells were fixed in 4% cold formaldehyde PBS for 15 minutes at room temperature, washed in PBS and incubated in blocking solution 5% BSA in PBS overnight at 4°C. S-phase cells were stained with a Click-iT EdU Cell Proliferation Assay kit according to the manufacturer’s instructions (ThermoFisher Scientific). Cells were then incubated with the indicated primary and Alexa Fluor 488 and 568 secondary antibodies in 2.5% BSA PBS solution for 1 hour at room temperature. Cover glasses were washed three times for 5 minutes each in between antibodies and mounted in ProLong Diamond with DAPI (ThermoFisher Scientific). Immunofluorescence images were captured using a Nikon Ni-E epifluorescent microscope under a 60X objective. Data were analysed using ImageJ. Signal for γH2AX was used as a marker of DNA breaks to outline the laser stripes, and the signal intensity in the channel of interest was then quantified. Nuclear background signal was subtracted from the intensity of the region of interest. Measurements were normalised to the earliest time-point DMSO control.

### Immunofluorescence

Cells were seeded on glass coverslips 24 hours before experiments. Cells were washed with PBS, fixed for 10 min with ice-cold 3.7% formaldehyde and washed three times in PBS. Cells were fixed for additional 5 min with ice-cold methanol. For pre-extraction, cells were washed once with ice-cold CSK buffer (10 mM Pipes pH 7.0, 100 mM NaCl, 300 mM sucrose, and 3 mM MgCl2), and pre-extracted twice for 2 min with ice-cold CSK containing 0.25% Triton X-100. After pre-extraction, cells were washed with ice-cold CSK buffer and fixed as described above. After rehydration in PBS, cells were blocked O/N with PBS containing 5% BSA. Coverslips were incubated for 1 h with primary antibodies in PBS/2.5% BSA and then washed five times with PBS and incubated with appropriate secondary antibodies coupled to Alexa Fluor 488 or 594 fluorophores in PBS/2.5% BSA. DNA was stained 10 μg/ml DAPI in PBS. After washes in PBS, coverslips were dipped in water and mounted on glass slides using ProLong™ Gold Antifade Mountant (ThermoFisher Scientific). Images of immunostained cells were acquired with an epifluorescent microscope (Nikon Ti-E) and foci analysis was performed using ImageJ automated counting. Representative images and co-localization analysis were acquired using a Zeiss LSM780 confocal microscope system.

#### Flow cytometry

Cells were harvested, washed with PBS and subsequently fixed in ice-cold methanol for 15 minutes at -20°C. After washing and rehydration in PBS containing 1% BSA, the cells were stained with 20 μg/ml of propidium iodide diluted in PBS 1% BSA and 10 μg/ml RNase A for 30 minutes at RT. For EdU analysis, cells were incubated with 10 μM EdU before harvesting. EdU was detected with a Click-iT EdU Cell Proliferation Assay kit (ThermoFisher Scientific). Cells were analyzed on a BD FACSCalibur™ flow cytometer. A minimum of 10,000 events was counted. Data analysis was performed using FlowJo.

#### Cellular fractionation

HEK293 cells were incubated in 2x volumes of Buffer A (10mM Hepes pH 7.4, 10mM KCl, 340mM sucrose, 10% glycerol, 2mM EDTA, 10mM NEM; protease and phosphatase inhibitors and 0.1% Triton X-100) on ice for 5 minutes. Samples were spun (500g, 3 min, 4°C) and the supernatants (cytosolic fraction) collected and stored. Nuclei were washed twice (500g, 3 min, 4°C) with Buffer A without Triton X-100 and burst in 2x volumes of hypotonic Buffer B (3 mM EDTA, 0.2 mM EGTA, 5 mM Hepes pH 7.9; 10 mM NEM; protease and phosphatase inhibitors) on ice for 10 min. After centrifugation at 1,700g for 3 min, the supernatant (nuclear soluble fraction) was collected and stored. The pellet (chromatin fraction) was washed twice (5,000 g for 5 minutes) with Buffer B. For chromatin nuclease extracts, chromatin fraction was washed in benzonase buffer (25 mM Tris pH 7.9; 25 mM NaCl; 2.5 mM KCl; 3 mM MgCl2) and incubated in the same buffer with 200U/ml benzonase on ice until DNA digestion was complete. Sample was spun at 20,000g for 5 min. The supernatant (chromatin soluble fraction) was quantified and analyzed by SDS-PAGE.

FA treatments were performed with 1 mM for 2 hours. Ubiquitination inhibitor (MLN7243, Chemietek) was typically added 15 minutes earlier and kept for the duration of FA treatment.

#### *In vitro* cleavage

SPRTN enzymatic reactions were performed as described previously (Vaz et al., 2016). Briefly, cleavage reactions were set up in 150mM NaCl and 25mM Tris (pH 7.4) in a PCR machine at 37°C. The reaction volume was typically 20μL. For proteolysis of SUMO-Topo-1: Recombinant Topo-1 (Abcam) was SUMOylated *in vitro* according to manufactur’s instructions (SUMOylation kit, Enzo Life Sciences), denatured at 95°C for 10 minutes, cooled on ice for 10 minutes, and incubated with recombinant SPRTN (1:1 ratio) for 16 hours at 37°C. For proteolysis of Topo-1 isolated from cells: YFP-Topo-1 was immunoprecipitated after 1 hour treatment with CPT (1μM) from HEK293 cells (80% confluence 15 cm dish) co-expressing HA-SUMO-1 using GFP-trap® beads (Chromotek). Briefly, cell extracts were prepared in denaturing conditions (1% SDS, 5 mM EDTA). Samples were diluted 10X in IP buffer (150 mM NaCl, 10 mM Tris, 0.5 mM EDTA pH 7.5) with 1% Triton and captured with GFP-trap beads for 2 hours at room temperature. Beads were washed 5 times in IP buffer and 2 times in reaction buffer (150 mM NaCl and 25 mM Tris pH 7.4). Beads containing denatured YFP-Topo-1 were divided in 3 and incubated with recombinant SPRTN (2 μg) for 16 hours at 37 °C. Reactions were analysed by Western blotting.

#### Co-immunoprecipitations

Cells were transfected with the plasmids of interest Lipofectamine® 3000 (ThermoFisher Scientific). Following treatment (according to the experiment), cells were washed twice with ice-cold PBS and lysed directly from the dish with ice-cold IP lysis buffer (50 mM Tris-HCl pH 7.4, 150 mM NaCl, 0.5% NP-40, protease and phosphatase inhibitors) containing 500 U/ml of benzonase. Flag-tag protein complexes were capture using the Anti-FLAG™ M2 Affinity Gel (Sigma-Aldrich) for 4 hours at 4°C. Beads were washed 5 times with IP lysis buffer and eluted using 3X Flag® peptide (Sigma-Aldrich). Flag-tag protein complexes were analysed by Western blotting.

#### DNA fiber combing

DNA fiber combing was performed according to GenomicVision instructions. Asynchronous HEK293 cells were labelled with 30 µM CldU (Sigma-Aldrich) for 30 minutes and then with 250 µM IdU (Sigma-Aldrich) for additional 30 minutes. Treatment with 450 µM FA was concomitant with IdU incubation. When specified, 50 µM 2-D08 (Sigma-Aldrich) or 5 μM MLN7243 (Chemietek) were added 15 minutes before CldU incubation and kept for the entire duration of the experiment; alternatively, DMSO was used. Cells were kept at 37°C for the length of the experiment. DNA replication was inhibited with 1x ice-cold PBS. DNA extraction and combing were performed with a FiberPrep® DNA extraction Kit (Genomic Vision) following manufacturer’s instructions. For fiber staining, anti-BrdU (for CldU) (Abcam) and anti-BrdU (for IdU) (BD Biosciences) were used. Anti-rat Cy5 (Abcam) and anti-mouse Cy3.5 Ab6946 were the respective secondary antibodies. Coverslips were scanned on a Genomic Vision FiberVision® platform. Quantification of IdU-labelled DNA tract lengths was done with FiberStudio® v0.15 software on at least 170 unidirectional fibers/condition. Resulting “Tract length” values (Kb) were represented in box and whiskers plots showing the median (horizontal band) with the 1^st^ and 3^rd^ quartile range (box) (the dots indicate the outliers) and statistically analysed using GraphPad Prism (two-tailed Mann-Whitney test).

#### Colony forming assay

HeLa cells were seeded at low density in 6-well plates and incubated overnight. Cells were exposed to increasing doses of FA diluted in cold DMEM for 20 minutes at 4°C, washed in warm PBS and incubated in fresh warm medium at 37°C for 7-10 days. After fixation and staining (1x PBS, 1% methanol, 1% Formaldehyde, 0.05% crystal violet), the number of clones was counted using an automated colony counter GelCount™ (Oxford Optronix). The number of colonies in treated samples was expressed as a percentage of colony numbers in the untreated samples.

#### Sister chromatid exchange assay

HeLa cells were grown in presence of 15 μM BrdU for 48 hours. After this incubation time, cells were treated for 10 minutes with 450 μM FA and DMSO or 10 μM 2-D08. After 2 washes with 1x PBS, cells were incubated for 16 hours at 37°C with medium containing 15 μM BrdU, 30 ng/ml colcemid solution, DMSO or 10μM 2-D08. Cells were collected, resuspended in pre-warmed optimal hypotonic solution (Genial Helix) and incubated at 37°C for 20 minutes. Cells were fixed with methanol:acetic acid (3:1, v/v). Staining was performed according to Clare, 2012.

### Quantification and statistical analysis

All experiments were reproduced at least two times, with similar results. The statistical method used for comparison between experimental groups was an unpaired t-test carried out using GraphPad Prism8. Statistical significance was expressed as a P value, where P<0.05 was considered a statistically significant difference.

### Data and code availability

This study did not generate datasets.

## KEY RESOURCES TABLE

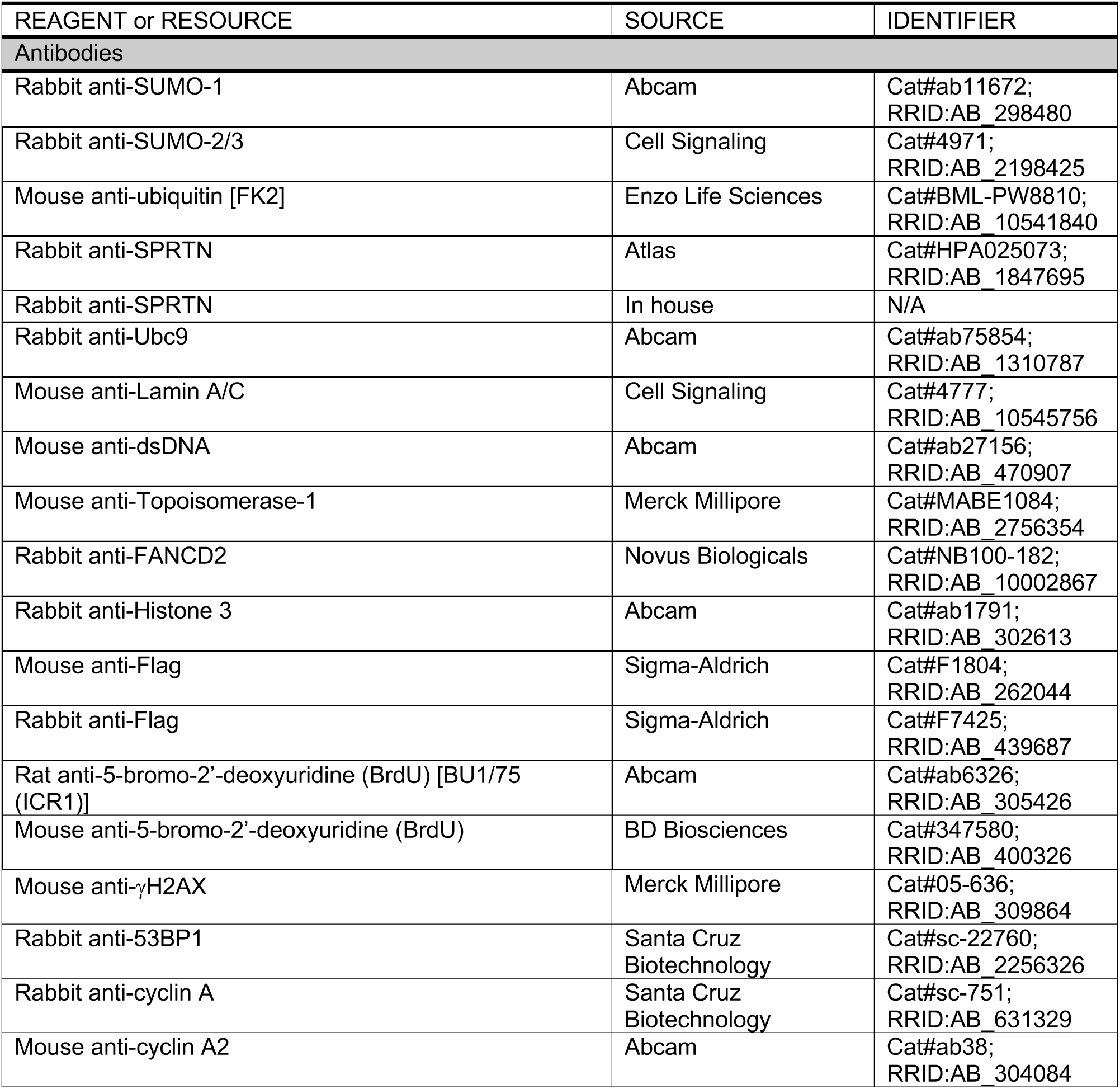

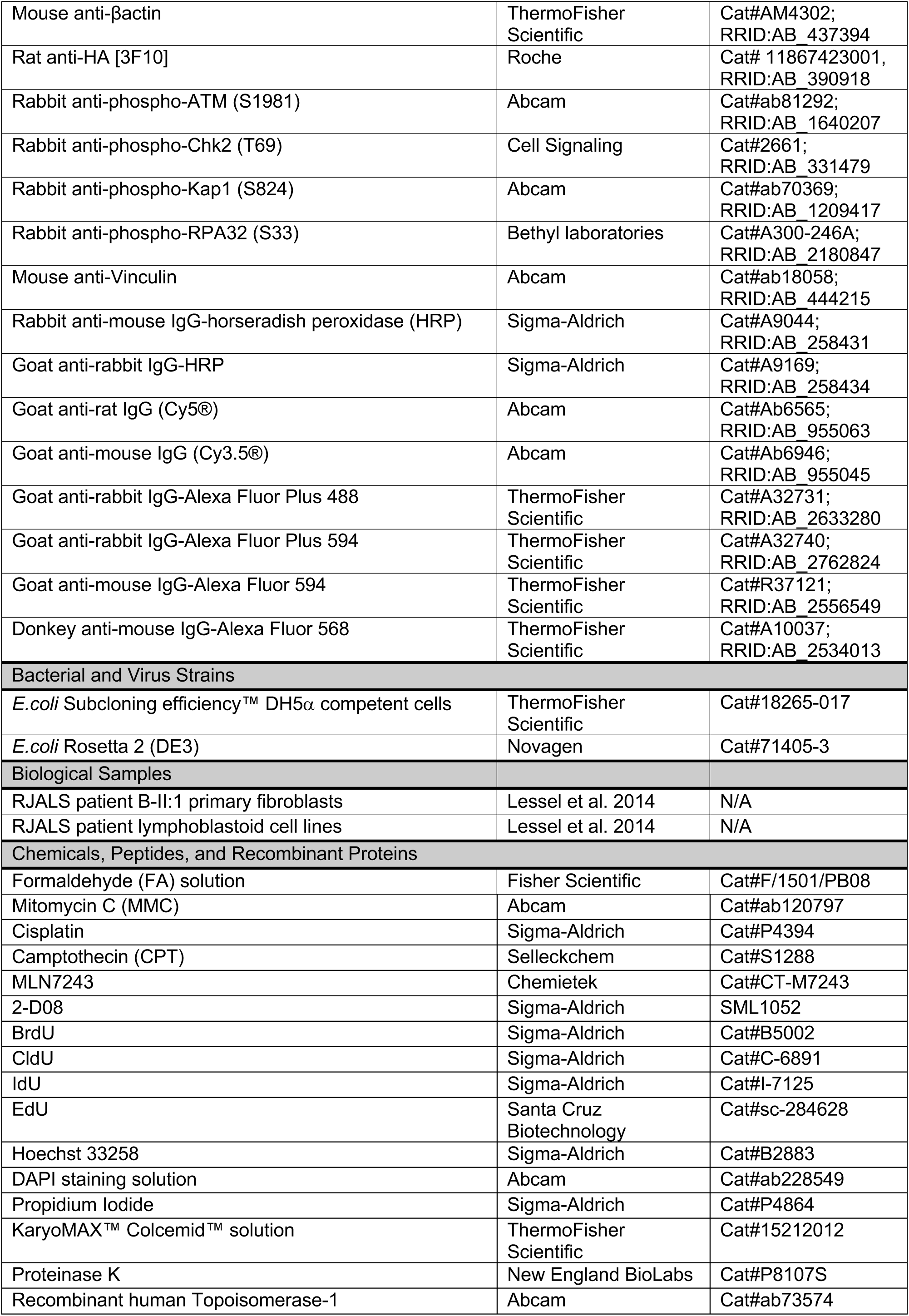

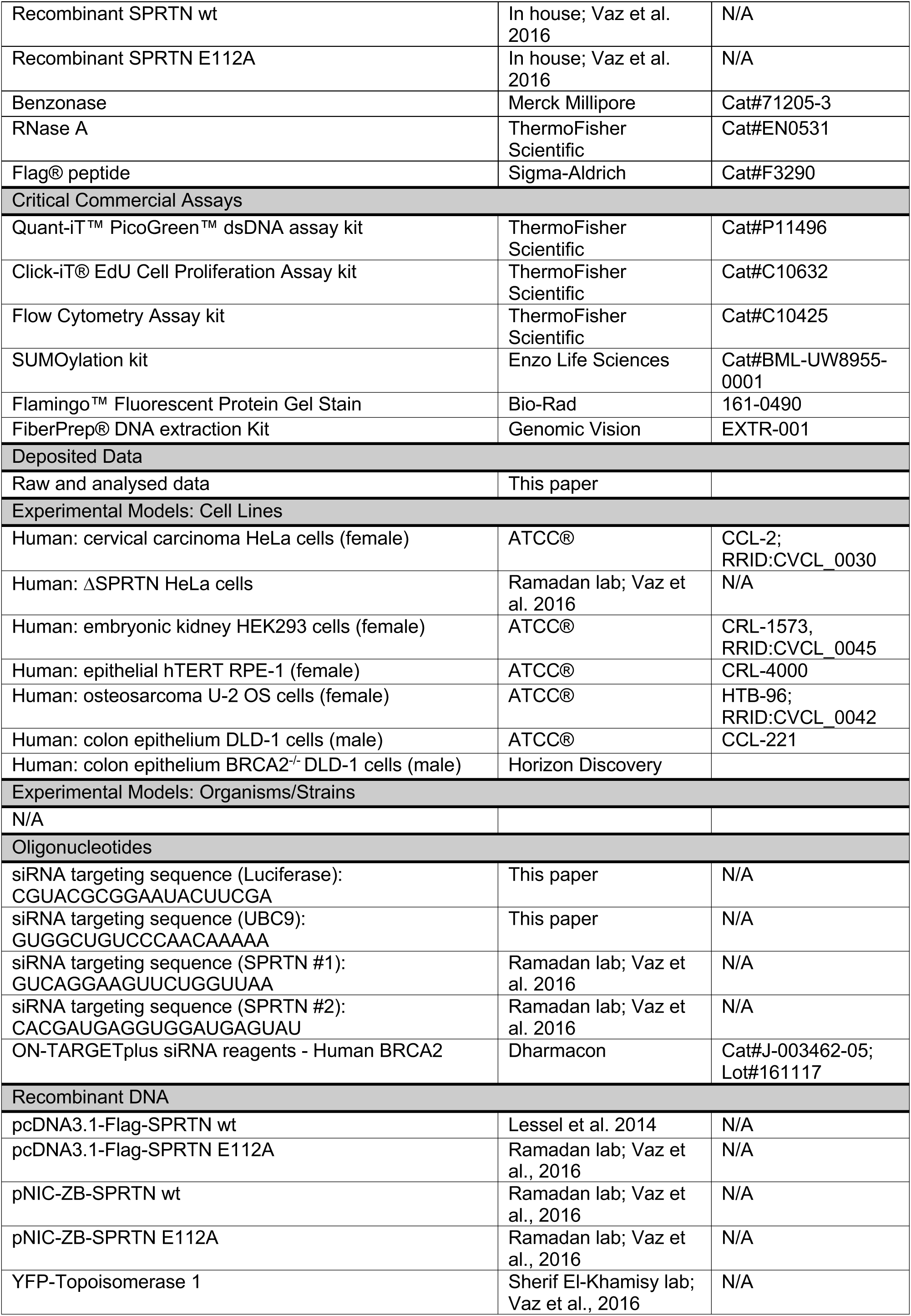

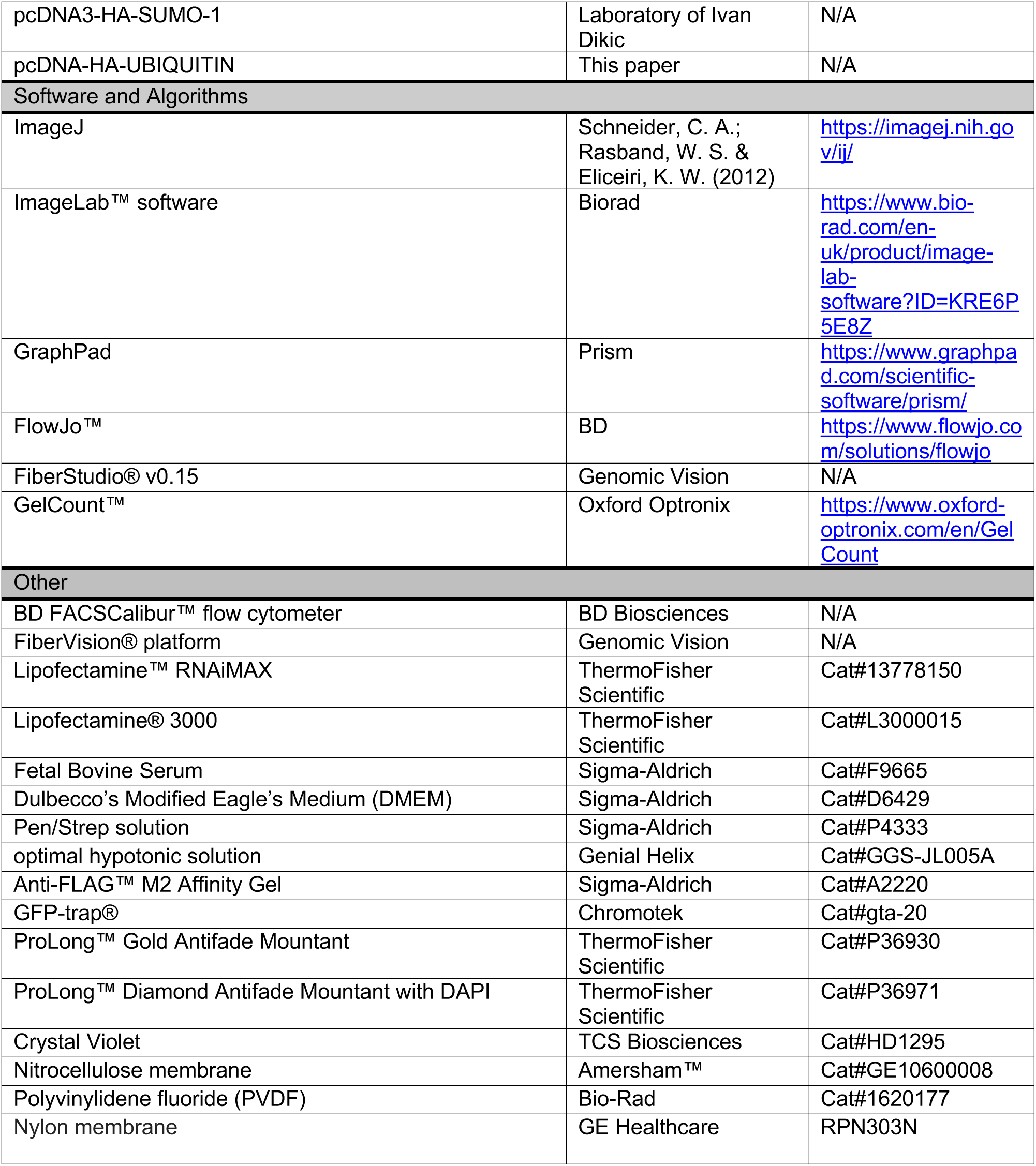

## REFERENCES

Andersen, M.E., Clewell, H.J., Bermudez, E., Dodd, D.E., Willson, G.A., Campbell, J.L., and Thomas, R.S. (2010). Formaldehyde: Integrating dosimetry, cytotoxicity, and genomics to understand dose-dependent transitions for an endogenous compound. Toxicol. Sci. 118, 716– 731.

Aparicio, T., Baer, R., Gottesman, M., and Gautier, J. (2016). MRN, CtIP, and BRCA1 mediate repair of topoisomerase II-DNA adducts. J. Cell Biol. 212, 399–408.

Ashour, M.E., Atteya, R., and El-Khamisy, S.F. (2015). Topoisomerase-mediated chromosomal break repair: An emerging player in many games. Nat. Rev. Cancer 15, 137–151.

Borgermann, N., Ackermann, L., Schwertman, P., Hendriks, I.A., Thijssen, K., Liu, J.C., Lans, H., Nielsen, M.L., and Mailand, N. (2019). SUMO ylation promotes protective responses to DNA -protein crosslinks . EMBO J. 38.

Branzei, D., Sollier, J., Liberi, G., Zhao, X., Maeda, D., Seki, M., Enomoto, T., Ohta, K., and Foiani, M. (2006). Ubc9- and Mms21-Mediated Sumoylation Counteracts Recombinogenic Events at Damaged Replication Forks. Cell 127, 509–522.

Ceccaldi, R., Sarangi, P., and D’Andrea, A.D. (2016). The Fanconi anaemia pathway: New players and new functions. Nat. Rev. Mol. Cell Biol. 17, 337–349.

Centore, R.C., Yazinski, S.A., Tse, A., and Zou, L. (2012). Spartan/C1orf124, a Reader of PCNA Ubiquitylation and a Regulator of UV-Induced DNA Damage Response. Mol. Cell 46, 625–635.

Clare, G. (2012). The In Vitro Mammalian Chromosome Aberration Test. In Genetic Toxicology: Principles and Methods J.M. Parry, and E.M. Parry, eds. (New York, NY: Springer New York), pp. 69–91.

Davis, E.J., Lachaud, C., Appleton, P., MacArtney, T.J., Näthke, I., and Rouse, J. (2012). DVC1 (C1orf124) recruits the p97 protein segregase to sites of DNA damage. Nat. Struct. Mol. Biol. 19, 1093–1100.

Desai, S.D., Liu, L.F., Vazquez-Abad, D., and D’Arpa, P. (1997). Ubiquitin-dependent destruction of topoisomerase I is stimulated by the antitumor drug camptothecin. J. Biol. Chem. 272, 24159–24164.

Fielden, J., Ruggiano, A., Popović, M., and Ramadan, K. (2018). DNA protein crosslink proteolysis repair: From yeast to premature ageing and cancer in humans. DNA Repair (Amst). 71, 198–204.

Flynn, J.M., Neher, S.B., Kim, Y.I., Sauer, R.T., and Baker, T.A. (2003). Proteomic discovery of cellular substrates of the ClpXP protease reveals five classes of ClpX-recognition signals. Mol. Cell 11, 671–683.

Ghosal, G., Leung, J.W.C., Nair, B.C., Fong, K.W., and Chen, J. (2012). Proliferating Cell Nuclear Antigen (PCNA)-binding protein C1orf124 is a regulator of translesion synthesis. J. Biol. Chem. 287, 34225–34233.

Gómez-Herreros, F., Schuurs-Hoeijmakers, J.H.M., McCormack, M., Greally, M.T., Rulten, S., Romero-Granados, R., Counihan, T.J., Chaila, E., Conroy, J., Ennis, S., et al. (2014). TDP2 protects transcription from abortive topoisomerase activity and is required for normal neural function. Nat. Genet. 46, 516–521.

Halder, S., Torrecilla, I., Burkhalter, M.D., Popović, M., Fielden, J., Vaz, B., Oehler, J., Pilger, D., Lessel, D., Wiseman, K., et al. (2019). SPRTN protease and checkpoint kinase 1 cross-activation loop safeguards DNA replication. Nat. Commun. 10.

Harper, J. V (2005). Synchronization of Cell Populations in G1/S and G2/M Phases of the Cell Cycle. In Cell Cycle Control: Mechanisms and Protocols, T. Humphrey, and G. Brooks, eds. (Totowa, NJ: Humana Press), pp. 157–166.

Henderson, C.J., and Wolf, C.R. (1992). Immunodetection of Proteins by Western Blotting. In Immunochemical Protocols, M.M. Manson, ed. (Totowa, NJ: Humana Press), pp. 221–233.

Hoa, N.N., Shimizu, T., Zhou, Z.W., Wang, Z.Q., Deshpande, R.A., Paull, T.T., Akter, S., Tsuda, M., Furuta, R., Tsusui, K., et al. (2016). Mre11 Is Essential for the Removal of Lethal Topoisomerase 2 Covalent Cleavage Complexes. Mol. Cell 64, 580–592.

Ide, H., Nakano, T., Salem, A.M.H., and Shoulkamy, M.I. (2018). DNA–protein cross-links: Formidable challenges to maintaining genome integrity. DNA Repair (Amst). 71, 190–197.

Interthal, H., and Champoux, J.J. (2011). Effects of DNA and protein size on substrate cleavage by human tyrosyl-DNA phosphodiesterase 1. Biochem. J. 436, 559–566.

Kiianitsa, K., and Maizels, N. (2013). A rapid and sensitive assay for DNA-protein covalent complexes in living cells. Nucleic Acids Res. 41, 1–7.

Larsen, N.B., Gao, A.O., Sparks, J.L., Gallina, I., Wu, R.A., Mann, M., Räschle, M., Walter, J.C., and Duxin, J.P. (2019). Replication-Coupled DNA-Protein Crosslink Repair by SPRTN and the Proteasome in Xenopus Egg Extracts. Mol. Cell 73, 574–588.e7.

Lessel, D., Vaz, B., Halder, S., Lockhart, P.J., Marinovic-Terzic, I., Lopez-Mosqueda, J., Philipp, M., Sim, J.C.H., Smith, K.R., Oehler, J., et al. (2014). Mutations in SPRTN cause early onset hepatocellular carcinoma, genomic instability and progeroid features. Nat. Genet. 46, 1239–1244.

Lin, C.P., Ban, Y., Lyu, Y.L., Desai, S.D., and Liu, L.F. (2008). A ubiquitin-proteasome pathway for the repair of topoisomerase I-DNA covalent complexes. J. Biol. Chem. 283, 21074–21083.

Link, A.J., and Labaer, J. (2011). Trichloroacetic acid (TCA) precipitation of proteins. Cold Spring Harb. Protoc. 6, 993–994.

Liu, P., Carvalho, C.M.B., Hastings, P.J., and Lupski, J.R. (2012). Mechanisms for recurrent and complex human genomic rearrangements. Curr. Opin. Genet. Dev. 22, 211–220.

Lopez-Mosqueda, J., Maddi, K., Prgomet, S., Kalayil, S., Marinovic-Terzic, I., Terzic, J., and Dikic, I. (2016). SPRTN is a mammalian DNA-binding metalloprotease that resolves DNA-protein crosslinks. Elife 5, 1–19.

Machida, Y., Kim, M.S., and Machida, Y.J. (2012). Spartan/C1orf124 is important to prevent UV-induced mutagenesis. Cell Cycle 11, 3395–3402.

Mao, Y., Desai, S.D., Ting, C.Y., Hwang, J., and Liu, L.F. (2001). 26 S Proteasome-mediated Degradation of Topoisomerase II Cleavable Complexes. J. Biol. Chem. 276, 40652–40658.

Maskey, R.S., Kim, M.S., Baker, D.J., Childs, B., Malureanu, L.A., Jeganathan, K.B., Machida, Y., Van Deursen, J.M., and Machida, Y.J. (2014). Spartan deficiency causes genomic instability and progeroid phenotypes. Nat. Commun. 5, 1–12.

Maskey, R.S., Flatten, K.S., Sieben, C.J., Peterson, K.L., Baker, D.J., Nam, H.J., Kim, M.S., Smyrk, T.C., Kojima, Y., Machida, Y., et al. (2017). Spartan deficiency causes accumulation of Topoisomerase 1 cleavage complexes and tumorigenesis. Nucleic Acids Res. 45, 4564–4576.

Mimnaugh, E.G., Chen, H.Y., Davie, J.R., Cells, J.E., and Neckers, L. (1997). Rapid deubiquitination of nucleosomal histones in human tumor cells caused by proteasome inhibitors and stress response inducers: Effects on replication, transcription, translation, and the cellular stress response. Biochemistry 36, 14418–14429.

Mórocz, M., Zsigmond, E., Tóth, R., Zs Enyedi, M., Pintér, L., and Haracska, L. (2017). DNA-dependent protease activity of human Spartan facilitates replication of DNA-protein crosslink-containing DNA. Nucleic Acids Res. 45, 3172–3188.

Mosbech, A., Gibbs-Seymour, I., Kagias, K., Thorslund, T., Beli, P., Povlsen, L., Nielsen, S.V., Smedegaard, S., Sedgwick, G., Lukas, C., et al. (2012). DVC1 (C1orf124) is a DNA damage-targeting p97 adaptor that promotes ubiquitin-dependent responses to replication blocks. Nat. Struct. Mol. Biol. 19, 1084–1092.

Nakano, T., Morishita, S., Katafuchi, A., Matsubara, M., Horikawa, Y., Terato, H., Salem, A.M.H., Izumi, S., Pack, S.P., Makino, K., et al. (2007). Nucleotide Excision Repair and Homologous Recombination Systems Commit Differentially to the Repair of DNA-Protein Crosslinks. Mol. Cell 28, 147–158.

Nakano, T., Katafuchi, A., Matsubara, M., Terato, H., Tsuboi, T., Masuda, T., Tatsumoto, T., Pil Pack, S., Makino, K., Croteau, D.L., et al. (2009). Homologous recombination but not nucleotide excision repair plays a pivotal role in tolerance of DNA-protein cross-links in mammalian cells. J. Biol. Chem. 284, 27065–27076.

Pommier, Y., and Marchand, C. (2012). Interfacial inhibitors: Targeting macromolecular complexes. Nat. Rev. Drug Discov. 11, 25–36.

Schwertman, P., Bekker-Jensen, S., and Mailand, N. (2016). Regulation of DNA double-strand break repair by ubiquitin and ubiquitin-like modifiers. Nat. Rev. Mol. Cell Biol. 17, 379–394.

Sordet, O., Larochelle, S., Nicolas, E., Stevens, E. V, Zhang, C., Shokat, K.M., Fisher, R.P., and Pommier, Y. (2009). RNA polymerase II is Hyperphopshorylated in Response to Topoisomerase I-DNA Cleavage Complexes and is Associated with transcription-and BRCA1-Dependent Degradation of Topoisomerase I. J. Mol. Biol 381, 540–549.

Sparks, J.L., Chistol, G., Gao, A.O., Räschle, M., Larsen, N.B., Mann, M., Duxin, J.P., and Walter, J.C. (2019). The CMG Helicase Bypasses DNA-Protein Cross-Links to Facilitate Their Repair. Cell 176, 167–181.e21.

Stingele, J., and Jentsch, S. (2015). DNA-protein crosslink repair. Nat. Rev. Mol. Cell Biol. 16, 455–460.

Stingele, J., Schwarz, M.S., Bloemeke, N., Wolf, P.G., and Jentsch, S. (2014). A DNA-dependent protease involved in DNA-protein crosslink repair. Cell 158, 327–338.

Stingele, J., Habermann, B., and Jentsch, S. (2015). DNA-protein crosslink repair: Proteases as DNA repair enzymes. Trends Biochem. Sci. 40, 67–71.

Stingele, J., Bellelli, R., Alte, F., Hewitt, G., Sarek, G., Maslen, S.L., Tsutakawa, S.E., Borg, A., Kjær, S., Tainer, J.A., et al. (2016). Mechanism and Regulation of DNA-Protein Crosslink Repair by the DNA-Dependent Metalloprotease SPRTN. Mol. Cell 64, 688–703.

Sun, Y., Jenkins, L.M.M., Su, Y.P., Nitiss, K.C., Nitiss, J.L., and Pommier, Y. (2019). A conserved SUMO-Ubiquitin pathway directed by RNF4/SLX5-SLX8 and PIAS4/SIZ1 drives proteasomal degradation of topoisomerase DNA-protein crosslinks. BioRxiv 707661.

Vaz, B., Popovic, M., Newman, J.A., Fielden, J., Aitkenhead, H., Halder, S., Singh, A.N., Vendrell, I., Fischer, R., Torrecilla, I., et al. (2016). Metalloprotease SPRTN/DVC1 Orchestrates Replication-Coupled DNA-Protein Crosslink Repair. Mol. Cell 64, 704–719.

Vaz, B., Popovic, M., and Ramadan, K. (2017). Review DNA – Protein Crosslink Proteolysis Repair. xx, 1–13.

Westphal, K., Langklotz, S., Thomanek, N., and Narberhaus, F. (2012). A trapping approach reveals novel substrates and physiological functions of the essential protease Ftsh in Escherichia coli. J. Biol. Chem. 287, 42962–42971.

