## Supplementary figures and images for "SPRTN protease and SUMOylation coordinate DNA-protein crosslink repair to prevent genome instability"

### Supplementary Figure 1

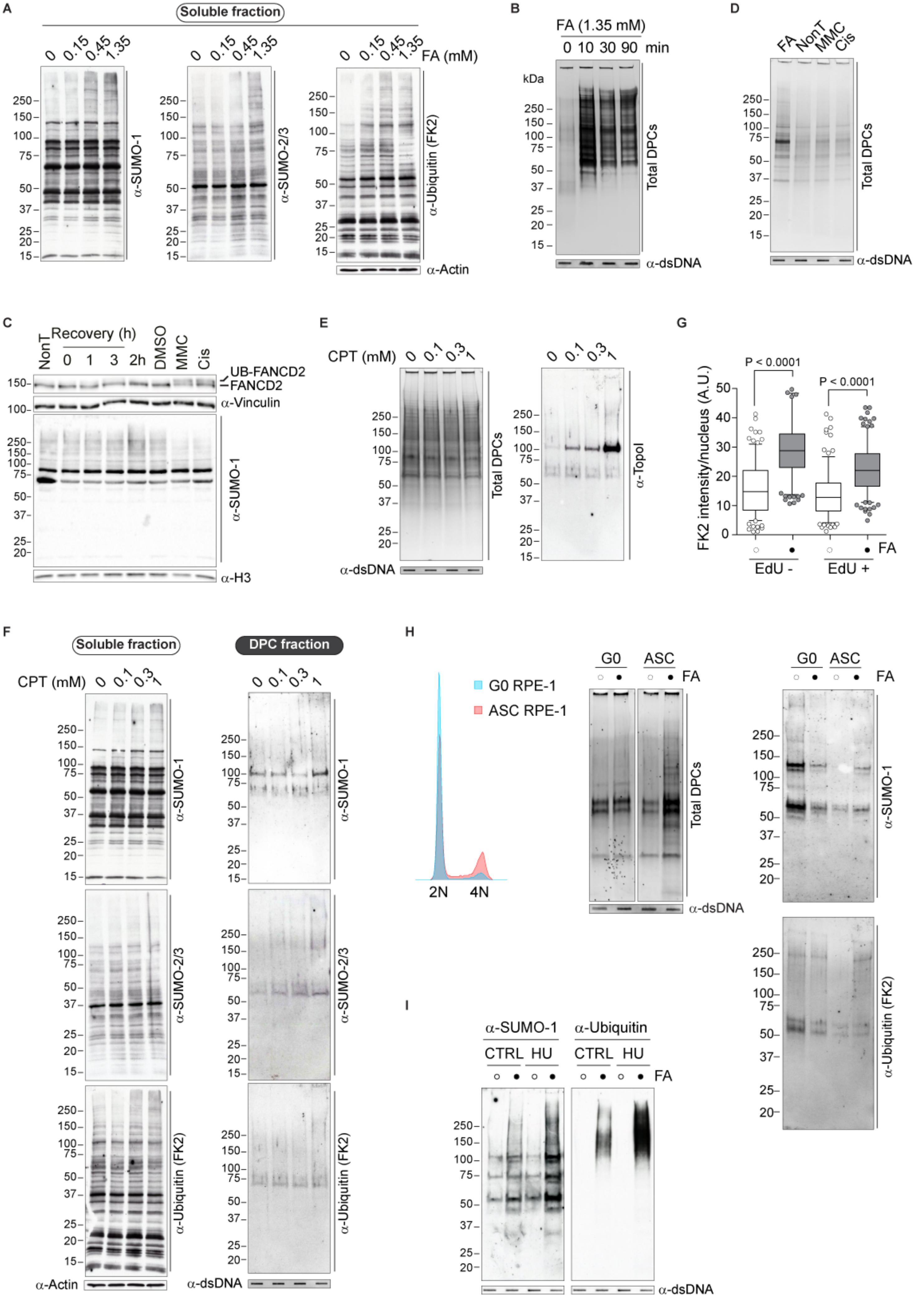

### Supplementary Figure 2

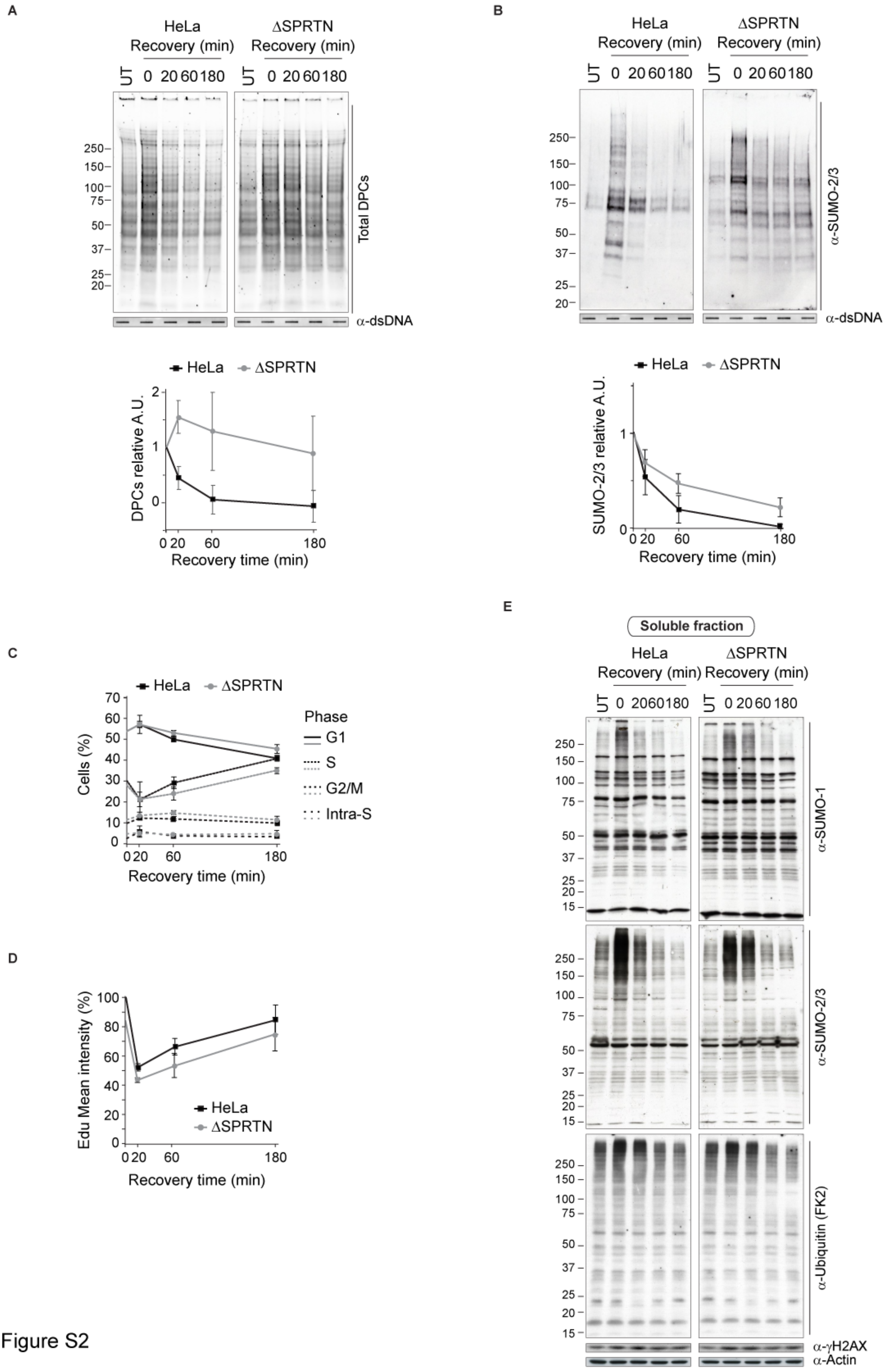

### Supplementary Figure 3

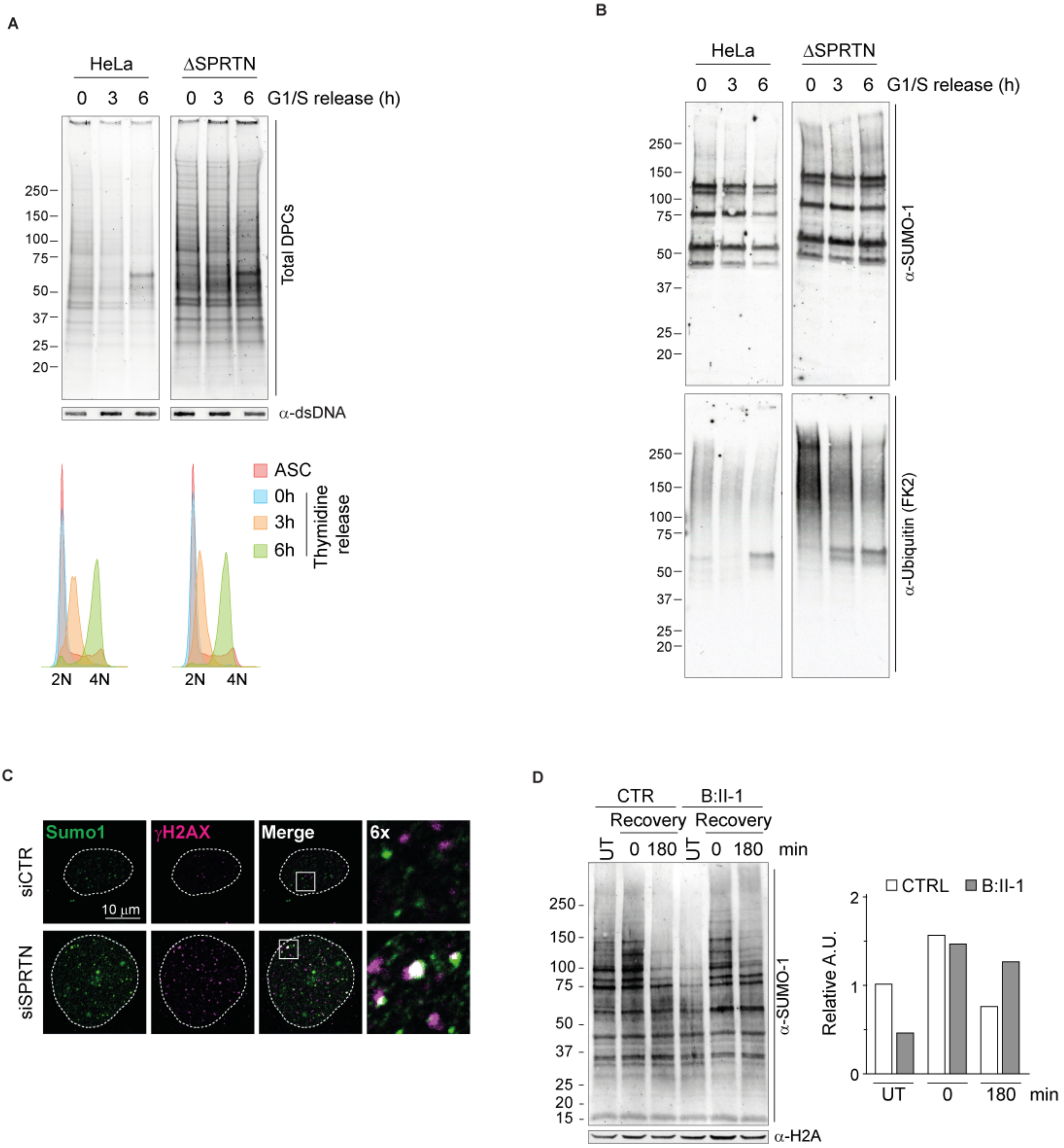

### Supplementary Figure 4

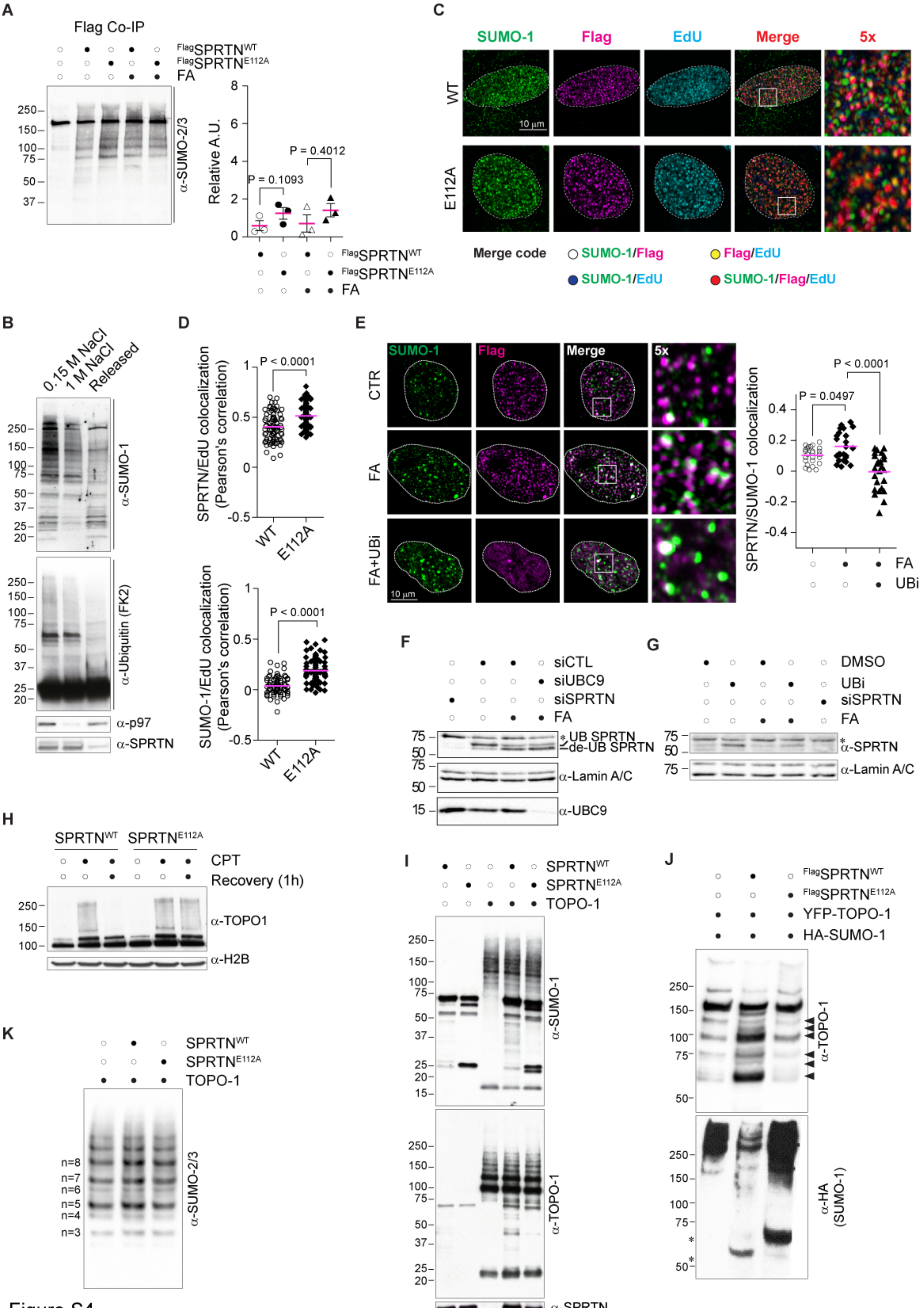

### Supplementary Figure 5

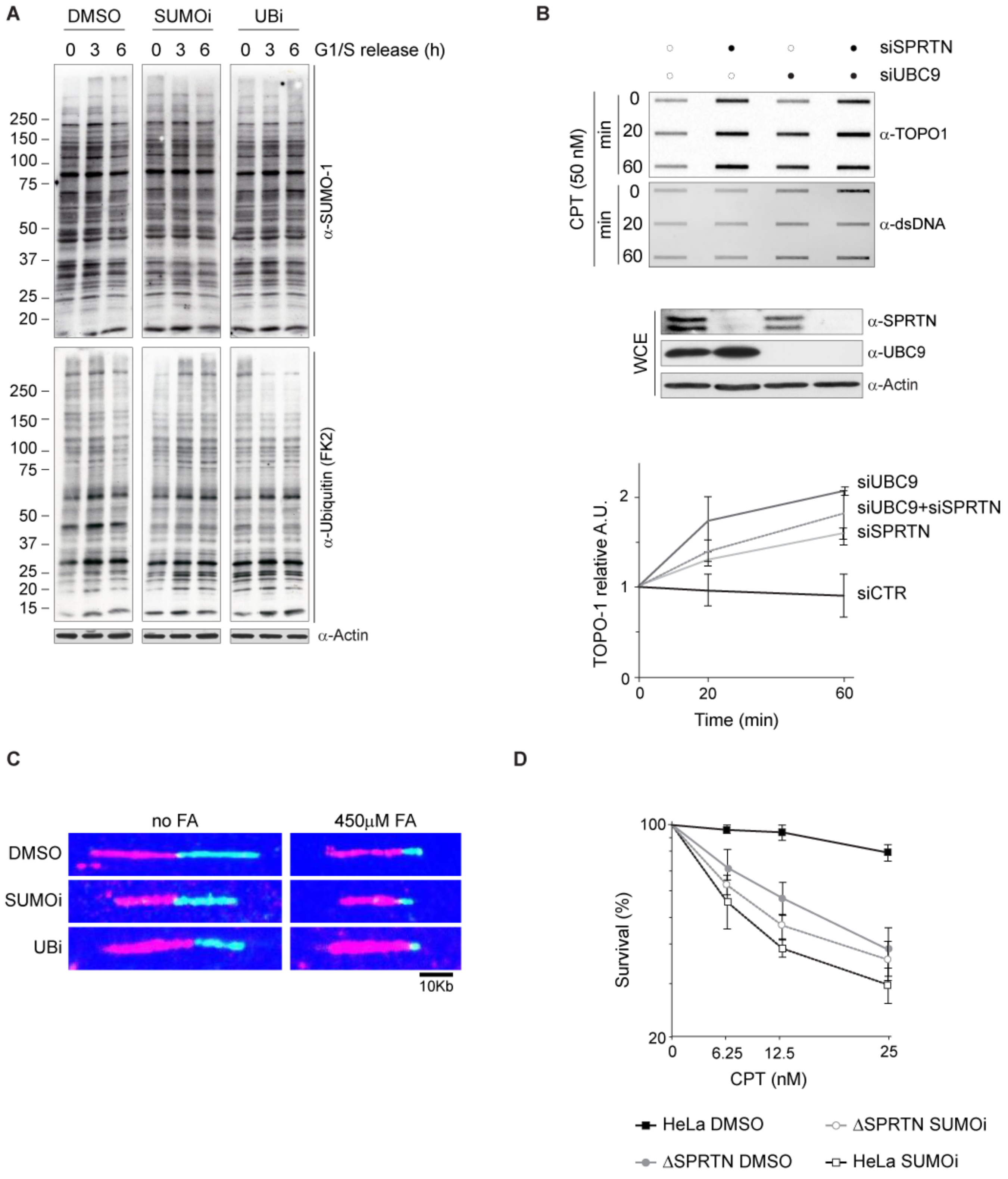

### Supplementary Figure 6

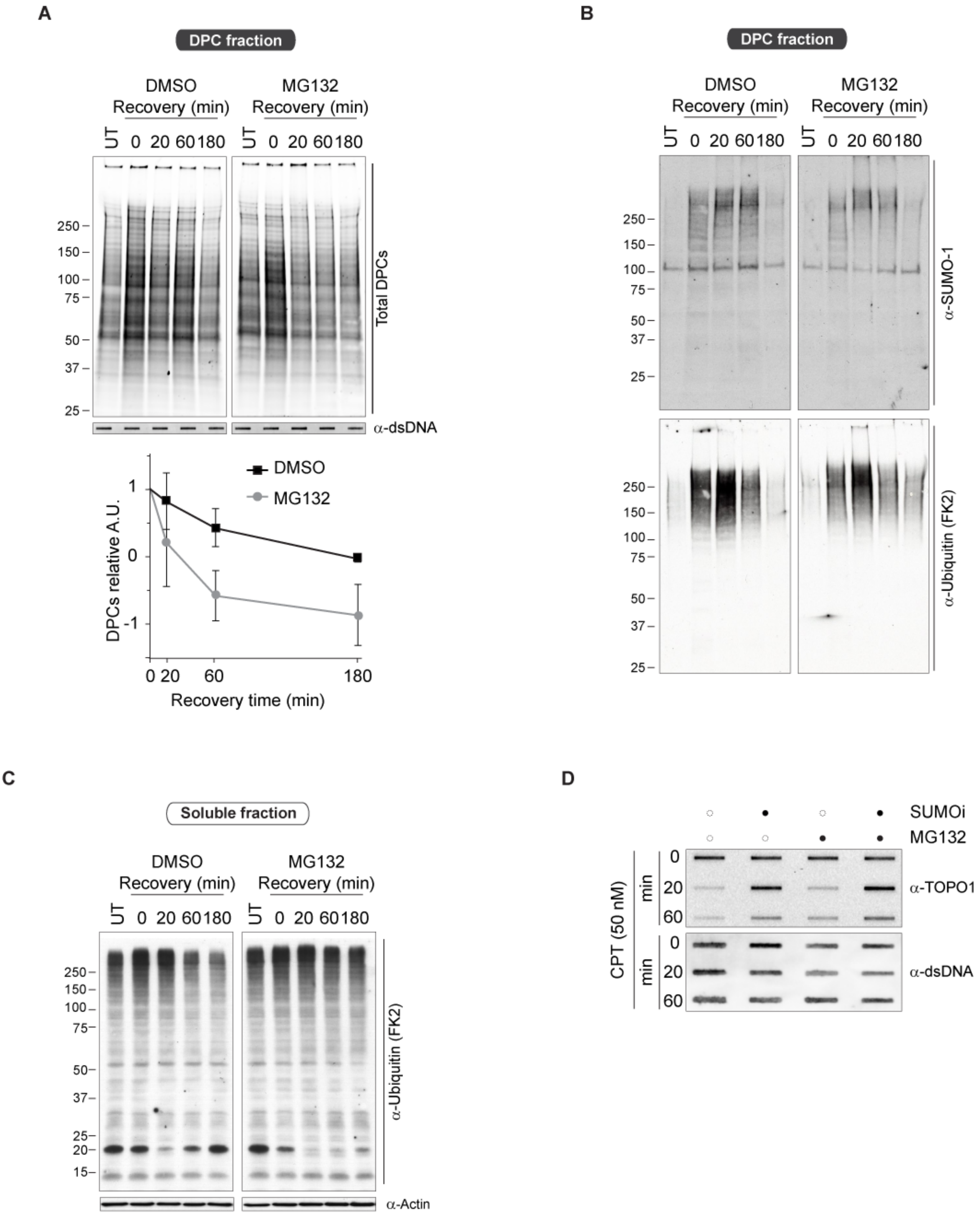

### Supplementary Figure 7

A

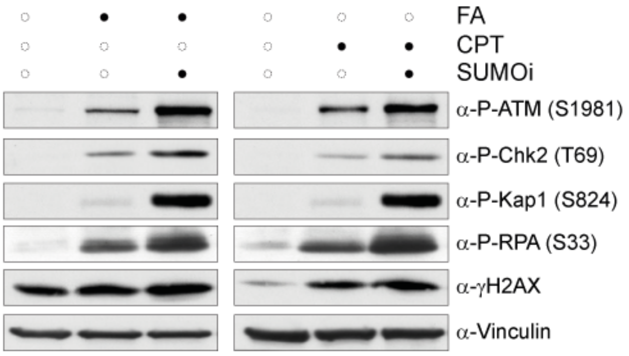


B

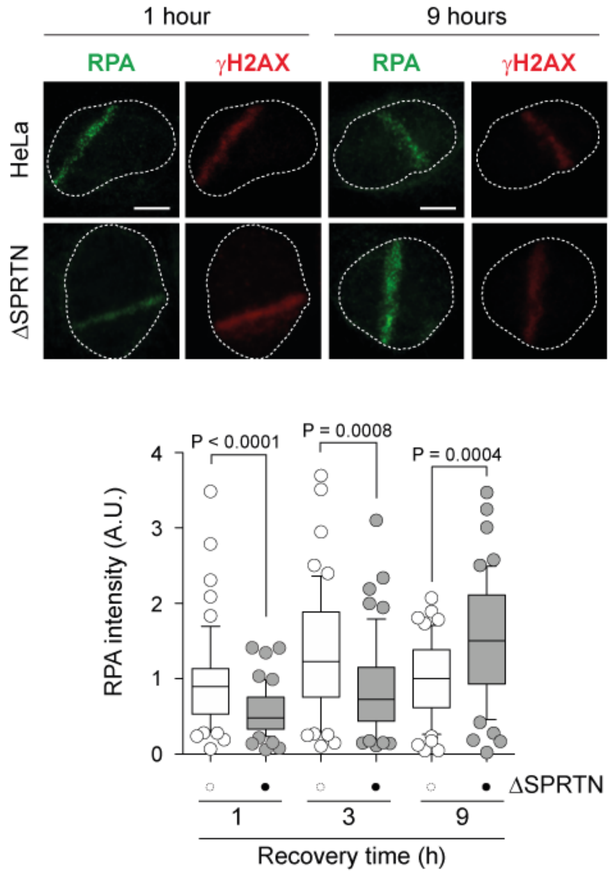


D

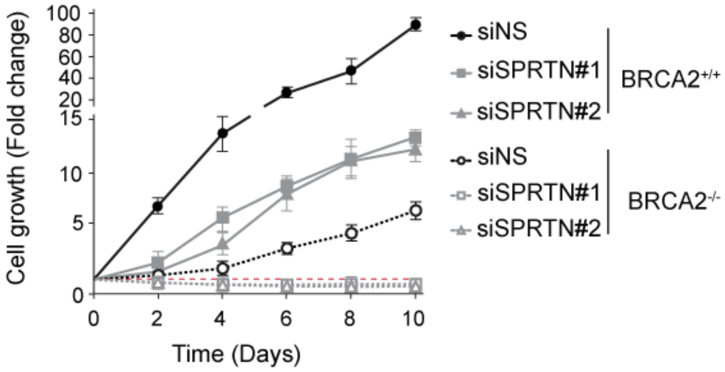


C

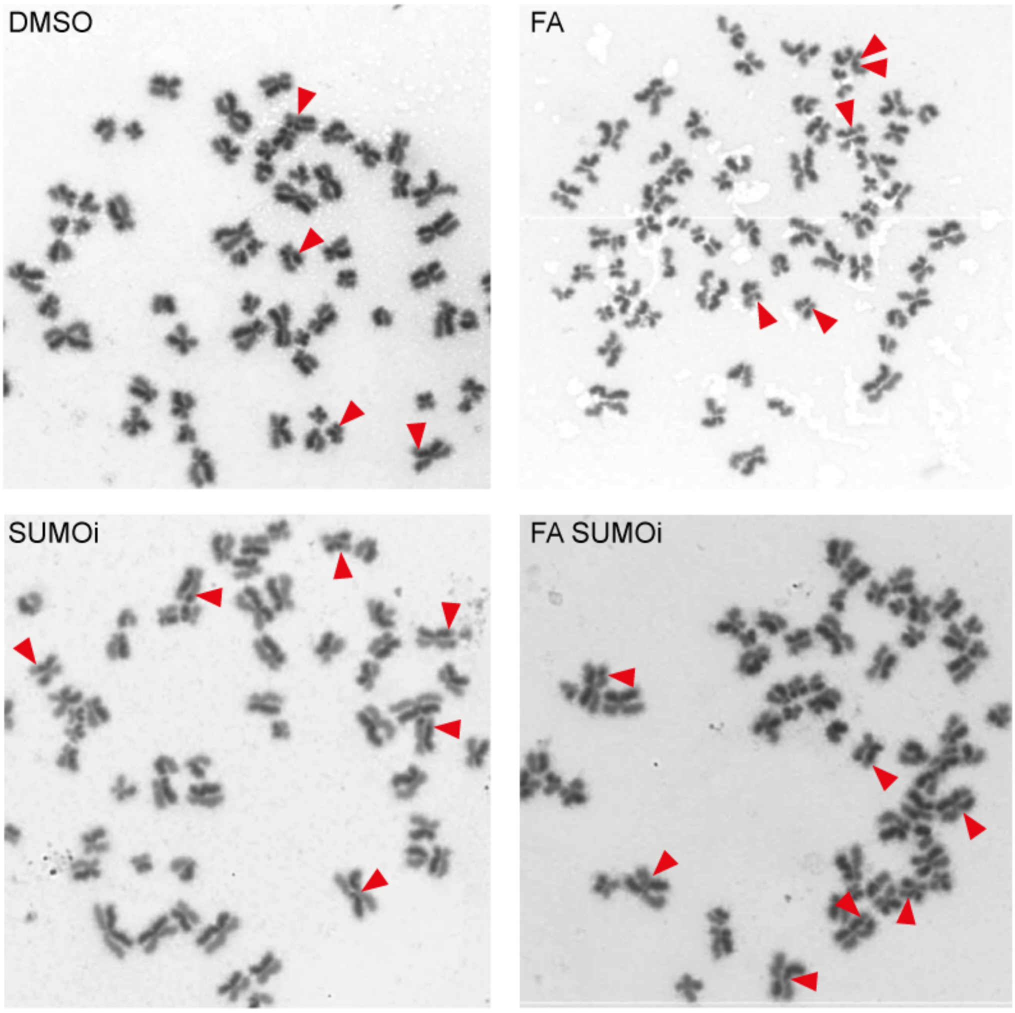


Figure S7
